# Proteasomal activity is differentially regulated in source and sink tissues of Arabidopsis

**DOI:** 10.1101/2022.11.16.516810

**Authors:** Haojie Wang, Joost T. van Dongen, Jos HM schippers

**Affiliations:** Department of Molecular Genetics, Leibniz Institute of Plant Genetics and Crop Plant Research (IPK) Gatersleben, 06466 Seeland, Germany; Institute of Biology I, Rheinisch-Westfälische Technische Hochschule Aachen University, 52074 Aachen, Germany

**Keywords:** Proteasome, Plant senescence, Proteostasis, Transcription Factors, Arabidopsis

## Abstract

Protein homeostasis controlled by the 26S proteasome plays a pivotal role in the adaption of plants to environmental stress, contributing to survival and longevity. During ageing in animals, proteasome activity declines resulting in senescence, however, in plants this is so far largely unexplored. Herein, we found that 26S proteasome capacity deteriorates with leaf age, while 20S proteasome activity increases. Interestingly, expression of proteasomal genes increases during leaf senescence, both at the steady-state mRNA level and poly-ribosome associated mRNA level. However, the increase in transcript level does not correlate with protein abundance and proteasome activity in senescing leaves. Furthermore, chemical inhibition of the proteasome results in accelerated leaf senescence. Interestingly, deterioration of proteasome activity in senescent leaves could be restored by cytokinin application. In Arabidopsis, feed-back regulation between proteasome activity and gene expression exists, and we propose that this is the cause for the high amount of proteasomal subunit mRNA during leaf senescence. In sink tissues like mature siliques and seeds, an increased 26S proteasome activity is observed. This increased activity is mainly due to enhanced proteasome assembly. This work provides new insights into the regulation of proteasome activity which deepens our understanding on source-sink relations and their impact on plant yield.

## Introduction

Plant senescence has been considered as an evolutionarily acquired developmental strategy that supports plant reproduction and survival (Schippers et al. 2015). During senescence, the plant proteome is actively reshaped, resulting in a massive turn-over of proteins. Loss of protein homeostasis is known as a hallmark of senescence in the worm *Caenorhabditis elegans*, resulting in ageing and death (Santra et al. 2019). The maintenance of protein homeostasis is controlled by the *de novo* protein synthesis through translation and targeted protein degradation. Considering the decline of protein synthesis during senescence, protein degradation might play an essential and leading role in plant ageing process and the progression of senescence (Lamattina et al. 1985; Hildebrandt et al. 2015).

Protein turn-over is largely controlled by the ubiquitin–proteasome system (UPS) and autophagy pathways. The UPS system typically removes single proteins that have been labelled with ubiquitin, which serves as degradation signal. In contrast, autophagy facilitates the bulk degradation of protein complexes, protein aggregates, cytoplasm, and organelles by the vacuole (Schreiber and Peter, 2014). Whereas autophagy only operates in the cytosol, the UPS system is also active in the nucleus. The highly selective degradation of single proteins by the UPS system is mainly to restrict the action of regulatory proteins, which are often short-lived as they control cellular differentiation or signaling processes (Xu and Xue, 2019). Recognized proteins are unfolded in an ATP-dependent manner and degraded by the proteasome. The 26S proteasome is a 2.5 MDa protease complex composed of two distinct sub-complexes (Vierstra et al. 2009), a 19S regulatory particle (RP) and a 20S core particle (CP). The RP consist of two sub-complexes (termed lid and base) and function as a selective filter of the CP, the proteolysis chamber. The lid is composed of ten regulatory particle non-ATPase (Rpn) subunits, while the base consists of six regulatory particle AAA-ATPase (Rpt) subunits and four Rpn subunits (Marshall and Vierstra, 2019). The Rpn subunits in the lid are mainly responsible for substrate recognition and deubiquitination, while the Rpt subunits in the base actively unfold substrates at the cost of ATP to present them to the CP (Tian et al 2011). The CP chamber is build-up of four stacked heteroheptameric rings of distinct α and β 20S proteasome subunits, whereby the proteolytic active sites are provided by PBA1, PBB1/2, and PBE1 in Arabidopsis. These three subunits have caspase-like, trypsin-like and chymotrypsin-like endopeptidase activities, respectively, enabling the proteasome to cleave most peptide bonds (Kurepa and Smalle, 2008). The assembly of active proteasome subunits is highly complex and requires numerous dedicated chaperones and maturation factors (Rousseau and Bertolotti, 2018). These include Ump1 chaperones that assist in the assembly of the 20S rings, and the PROTEASOME ACTIVATING PROTEIN 200 (PA200) chaperone that can transiently cap the 20S particle (Li et al., 2007; Dange et al., 2011). Still, the exact function of these proteins in the assembly of the 26S proteasome in plants is poorly understood.

For maintaining protein homeostasis, proteasome activity is versatilely regulated, ranging from the synthesis of its subunits, the rate of its assembly and disassembly, and post translational modifications (PTMs) to control its activity, substrate recognition and subcellular localization (Livneh et al. 2016, Wang and Schippers, 2019). Transcriptional regulation of proteasome subunits represents the first layer of control, whereby it has been shown that the genes encoding the subunits act as a regulon as they are simultaneously controlled by common upstream transcriptional regulators. In all eukaryotes analogous transcriptional regulators have been found, including the C2H2-type transcription factor Rpn5 in yeast (Mannhaupt et al. 1999), the basic leucine zipper transcription factor Nrf1 (Nuclear Factor Erythroid-derived 2-Related Factor 1) in mice and humans (Radhakrishnan et al. 2010; Radhakrishnan et al. 2014), and two NAM, ATAF1/2, and CUC2 (NAC) transcription factors ANAC053 and ANAC078 in Arabidopsis (Yabuta et al. 2011; Nguyen et al. 2013; Gladman et al. 2016). Apart from regulation at the transcriptional level, also regulation of proteasome assembly plays an important role (Marshall and Vierstra, 2019). At the moment, several chaperones have been identified in plants, including proteasome biogenesis-associated chaperone (PBAC) that likely promote CP assembly, and UMP1 chaperones that are responsible for connecting CP half-barrels (Gemperline et al. 2019). Mutants lacking UMP1 or PBAC chaperones are hypersensitive to proteotoxic stress, salt stress and osmotic stress (Gemperline et al. 2019; Wang et al. 2019). In animals it was shown that proteasome activity declines with age (Vilchez et al. 2014), however, in plants the proteasome is reported to be highly active in senescing leaves (Roberts et al. 2002; Poret et al. 2016). In addition, some studies suggest that proteasome regulation during ageing depends on the cell type being studied, like somatic and reproductive tissue (Fredriksson et al. 2012; Tsakiri et al. 2013). Furthermore, inhibition of the proteasome via specific inhibitors induce young animal cells to senesce or cause cell death (Chondrogianni et al. 2003; Harhouri et al 2017) indicating a key role of the proteasome in cell rejuvenation and survival (Vilchez et al., 2012).

Whereas the role of the proteasome in animals has been shown to involve the maintenance of stemness and prevent ageing and senescence (Vilchez et la., 2012; Vilchez et al., 2014), in plants this is not really clarified. In contrast to animals, plants are autotrophic, sessile and capable of resource allocation, and might therefore have a different regulation of ageing (Thomas, 2002). One divergent point is the relationship between sink and source tissues, which affect plant yield and ensure reproductive success (Schippers et al., 2015). As net importers of nutrients and assimilates, sink tissues receive precursors and building blocks from source tissues, including senescing tissues (Thomas, 2013; Schippers et al. 2015). Given that proteasome regulation during plant senescence has not been systematically studied, and considering the importance of protein homeostasis and proteasome functioning in plant development, the main purpose of this study was to offer a detailed characterization of proteasome functionality during plant senescence. Here, we found that 26S proteasome activity in leaves declines with age while the expression of proteasomal subunits is increased. Interestingly, the increased expression level of the subunits does not result in an increase in protein levels. In line with this, 26S proteasome activity declines with age, while 20S activity appears to increase. Interestingly, cytokinin treatment restores 26S proteasome activity. In contrast, in mature siliques and seeds, 26S proteasome activity is maintained, probably through enhanced proteasome assembly. Or study clearly indicates differential regulation of the proteasome in an age and tissue-dependent manner in Arabidopsis.

## Results

### During leaf senescence proteasomal activity shifts towards 20S complexes which is reversable by cytokinin

To understand the regulation of the proteasome during leaf ageing and senescence, proteasome activity was determined in young, mature and senescing first leaf pairs of Arabidopsis. For this, total protein was extracted and active proteasome complexes were resolved by native PAGE electrophoresis using a fluorogenic substrate (Suc-LLVY-AMC). The abundance of double-capped (DC) and single-capped (SC) 26S proteasome is the highest in young leaves, but drastically decreases with age, especially during leaf senescence (**Figure 1A-B**). In contrast, 20S proteasome activity is relatively low in young leaves, but is strongly induced in senescing leaves (**Figure 1A-B**). To further understand the impact of proteasome activity on the onset of leaf senescence, detached leaves were treated with either DMSO or the proteasome inhibitor MG132 and incubated in the dark (Buchanan-Wollaston et al., 2005). MG132 treatment accelerated chlorophyll loss in detached leaves as compared to DMSO treatment alone (**Figure 1C**). This implies that decreased proteasome activity promotes the induction of senescence and cell death.

**Figure 1.**
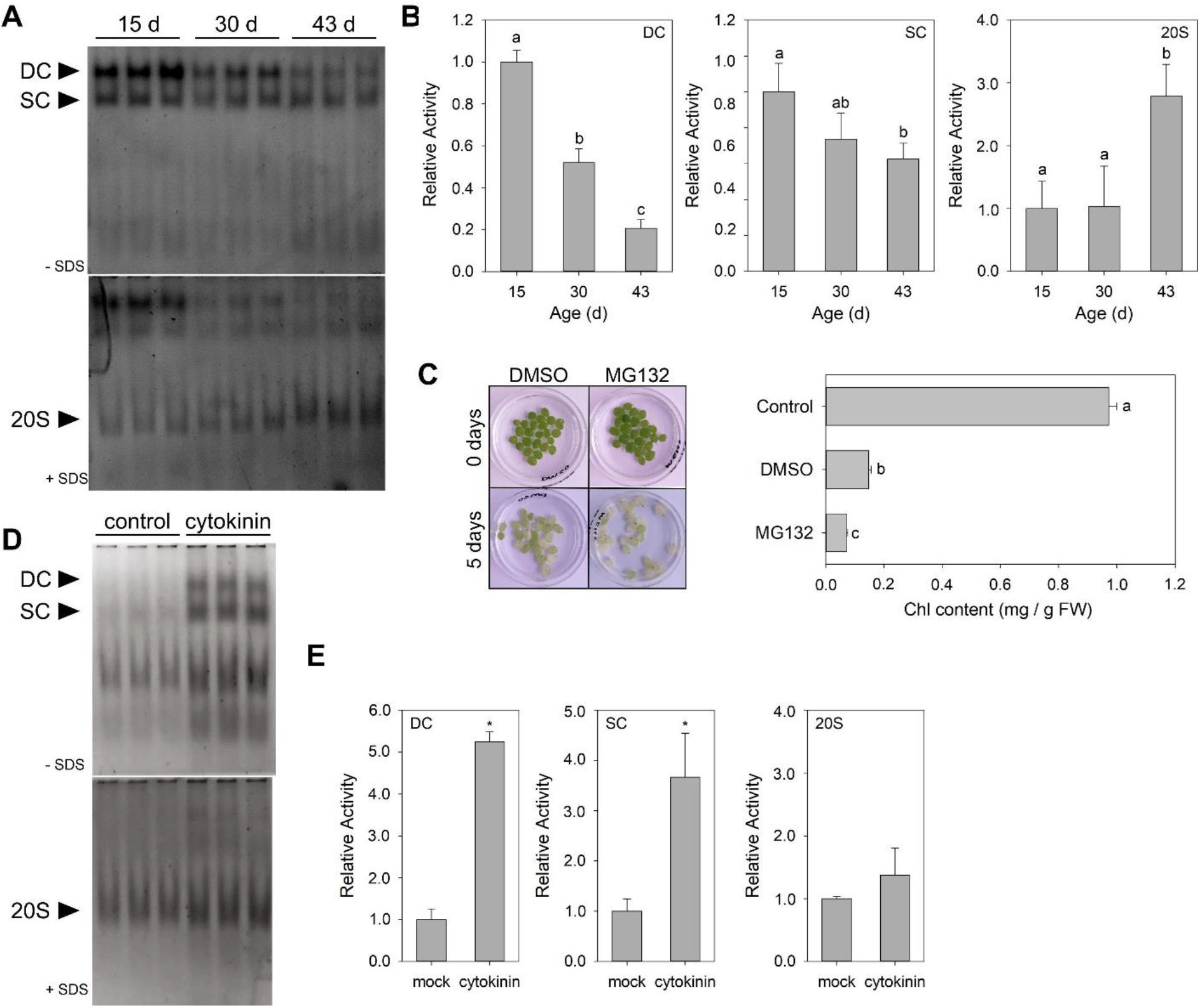
Proteasome activity declines with age but is induced by cytokinin. A, Suc-LLVY-AMC (100 μM) stain of total lysates resolved on 4% Native-PAGE showing the abundance and peptidase activity of different proteasome particles (DC: double-capped 26S proteasome; SC: single-capped 26S proteasome; 20S: 20S proteasome). Total protein from the first leaf pair was isolated at day 15 (young), day 30 (mature) and day 43 (senesced) and three independent biological replicates were loaded on gel. B, Bar graphs show relative activity for the different proteasomal complexes as based on digital quantification of band intensities from the activity gels. Data is expressed relative to the activity present at day 15, which was set at 1.0. Significant differences were determined using one-way ANOVA with post-hoc Tukey HSD Test. Different letters indicate significant differences (p < 0.05). C, Dark-induced leaf senescence was performed with first leaves isolated from 30-d-old Arabidopsis plants. Leaves were floated on water containing DMSO or 30μM MG132. Pictures show leaves before and after 5 days in darkness. After the treatment the chlorophyll content was determined. Bars indicate chlorophyll content and different letters indicate significant differences according to one-way ANOVA and post-hoc Tukey HSD Test (p < 0.05). D, Proteasomal activity gels (Suc-LLVY-AMC stained) of total lysates from cytokinin or mock treated plants, legend as in A. E, Bar graphs show relative activity for the different proteasomal complexes as based on quantification of band intensities. The activity after cytokinin treatment is expressed relative to the activity without, which was set at 1.0. Asterisks indicate significant differences, Student’s t test (p < 0.05).

The phytohormone cytokinin is well known for its ability to delay senescence or cause regreening in plants (Gan and Amasino, 1995; Guan et al. 2014). Therefore, we tested if cytokinin treatment might also impact on the formation of proteasomal complexes and activity. First of all, we found that cytokinin treatment slows down chlorophyll degradation of detached leaves under darkness and even counteracts the accelerated senescence induced by the proteasome inhibitor (**Supplemental Figure S1A**). To determine if cytokinin impacts on proteasomal complexes and activity, 30-day-old plants were sprayed with 80 μM cytokinin or mock treatment every day and the first leaf pair was collected after 6 days of treatment for protein isolation. Interestingly, cytokinin treatment promotes DC and SC 26S proteasome activity in the first leaf pair, while the 20S activity was hardly affected (**Figure 1D-E**). To determine if proteasomal subunit abundance was induced by cytokinin, we used the *pPAG1:PAG1-FLAG pag1-1* lines, which allow for the immunopurification of the whole 26S proteasome from crude plant extracts in the presence of ATP (Book et al., 2010). Immunopurification revealed that CP and RP subunits are more abundant after cytokinin treatment (**Supplemental Figure S1B**). In addition, western blot analysis of of RPN11 and PA200 confirmed that cytokinin increases their abundance (**Supplemental Figure S1C)**. Overall, our analysis reveals that 26S proteasome activity declines with leaf ageing and that cytokinin is capable of preventing this.

### Dynamic regulation of proteasomal gene expression and protein abundance during proteotoxic stress and leaf senescence

To better understand how 26S-proteasome abundance is controlled during leaf ageing and senescence, we generated expression reporters by cloning the promoter of several proteasomal subunits in front of the GUS gene. These include reporters for the genes encoding the proteasome chaperone *PA200*, and the RP particles *RPN6*, *RPN11*, *RPT1a*, *RPT1b*, *RPT3*, and *RPT5a*. To test whether these reporters are functional, plants were exposed to proteotoxic stress, as this was shown previously to induce the expression of proteasomal subunit genes (Gladman et al., 2016). Upon MG132 treatment, transgenic seedlings containing *pRPN11:GUS*, *pRPT1a:GUS*, *pRPT3:GUS*, *pPA200:GUS*, and *pRPT5a:GUS* showed an increased staining as compared to the control plants (**Figure 2A**). Still, plants carrying the *pRPT1b:GUS* and *pRPN6:GUS* construct did not respond to proteotoxic stress. To investigate if an increase in expression level is also followed by an increase in protein level, we used subunit specific antibodies for PA200, RPN11 and PBA1 and determined their abundance by western blot. The induction of proteotoxic stress by either MG132 or Bortezomib treatment results in an increased protein abundance for RPN11, PA200, and PBA1 (**Figure 2B**). Thus, the induction of transcript level by proteotoxic stress is translated into an increase in protein level of proteasomal subunits. Subsequently, the established promoter reporter lines were used to determine if during leaf senescence the expression of proteasomal subunits is altered. In contrast to the decrease in DC and SC 26 proteasome activity, the activity of the GUS reporter lines (*pPA200:GUS, pRPN6:GUS, pRPN11:GUS*, *pRPT1a:GUS*, and *pRPT5a:GUS*) showed an increase during leaf senescence (**Figure 2C**). These results imply that both proteotoxic stress and senescence promote the expression of subunit genes of the proteasome. To further asses the expression of proteasomal genes during leaf ageing, we generated a qPCR platform to assess the expression of all subunits at five different time-points during leaf growth (**Figure 2D**). The expression analysis revealed that most proteasomal subunits are induced during leaf senescence as compared to their expression level in young leaves (**Supplemental Dataset S1**). Contrary to the inhibitor treatment, the protein level of proteasomal subunits did not increase during leaf senescence (**Figure 2E**). Moreover, a decline for PA200 and RPN11 was observed at the onset of senescence. In addition, the 20S subunit PBA1 appeared to stay at a constant level. The protein level for the proteasomal subunits during leaf senescence parallels the detoriation of 26S proteasome activity during leaf ageing (**Figure 1A**).

**Figure 2.**
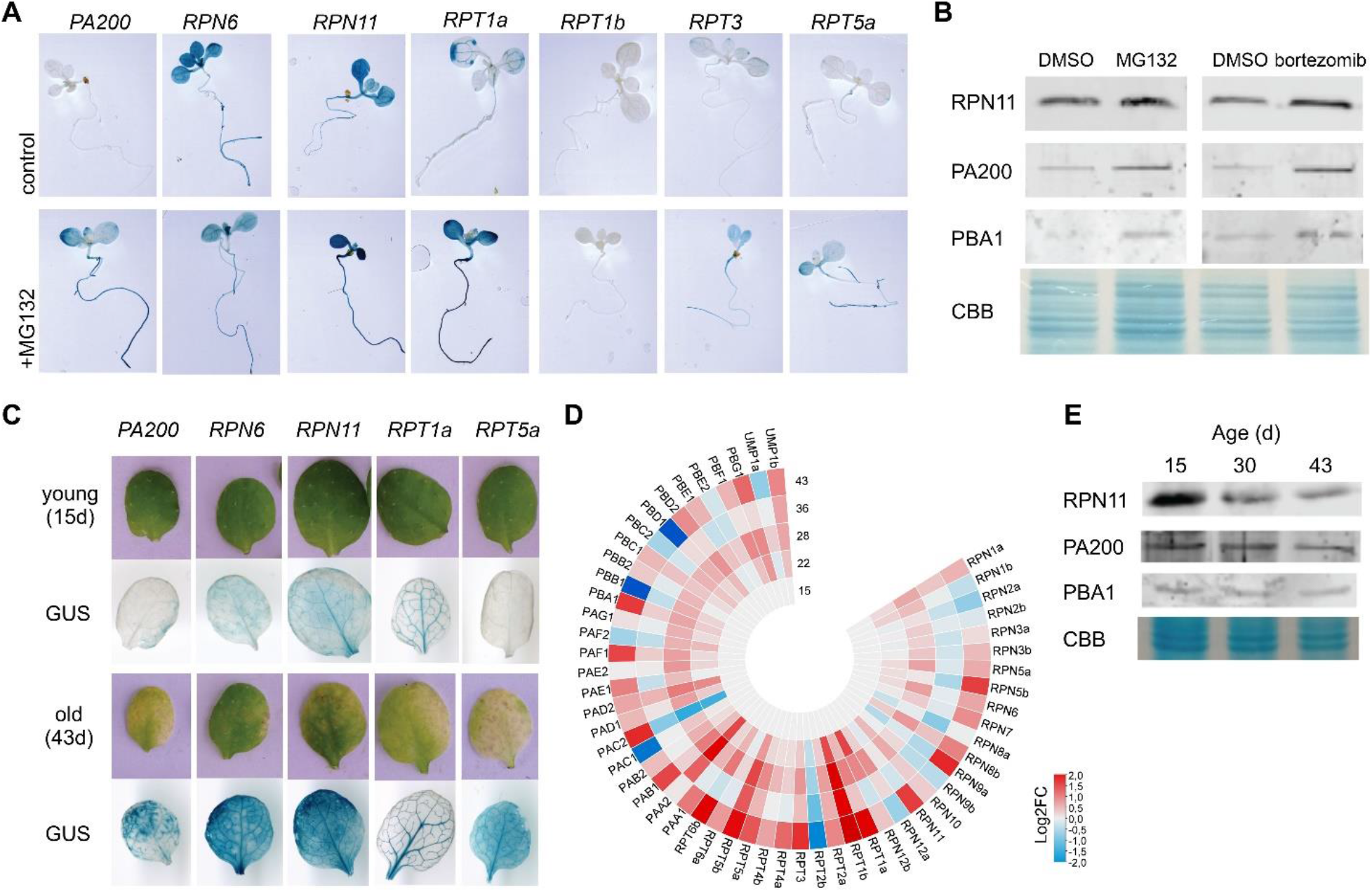
Proteasomal gene regulation during proteotoxic stress and leaf senescence. A, Effects of proteotoxic stress on the expression of proteasome promoter:GUS transgenes. Transgenic lines were grown in ½ strength Murashige and Skoog (MS) with/without 15μM MG132 for 10 days and then incubated overnight with the X-Gluc substrate. B, Proteasomal subunit abundance during proteotoxic stress. Col-0 seedling of 8 days were treated for for 2 days with either 30μM MG132, 2μM bortezomib or DMSO as a control. Total protein was separated and blotted and individual subunits were detected using specific antibodies. Coomassie Brilliant Blue (CBB) stained gels were used loading control. C, Expression of proteasomal subunit genes during leaf senescence as visualized by promoter:GUS transgenes. Shown are the results for the first leaf pair, harvested either from 15-d-old plants (young) or 43-d-old plants (old). D, A heat map summarizing the expression of proteasomal subunit genes during leaf ageing. The expression of proteasomal genes is shown for 5 time points, whereby the expression levels is given as compared to the level at day 15 (Log2FC). E, Proteasomal subunit abundance during leaf ageing. The first leaf pair of Col-0 was harvested at the given time point, and total protein was separated and blotted. The abundance for selected subunits was detected using specific antibodies. CBB stained gels were used loading control.

### Transcriptional regulation of proteasomal subunit genes during senescence

To understand the transcriptional regulation of proteasomal genes during senescence, we identified upstream transcription factors based on previously published DAP-seq data (O’Malley et al., 2016). We screened transcription factor families for members that are able to interact with nearly all proteasomal subunits genes (**Supplemental Dataset S2**), as controlling all subunits would enable the upstream regulator to modulate proteasomal activity. Amongst the NAC transcription factors, binding data for 50 members has been reported, and the known proteasomal regulators ANAC053 and ANAC078 where both found in the DAP-seq data to interact with nearly all genes encoding the proteasome (**Figure 3A; Supplemental Dataset S2**). Next to that, we found that ANAC010, ANAC046, ANAC055, ANAC071 and ANAC081 (ATAF2) were able to interact with nearly all proteasomal genes, suggesting that they might act in parallel to ANAC053 and ANAC078. Subsequently the expression of the NAC transcription factors that act upstream of the proteasomal genes during leaf ageing was analyzed. To this end, available temporal RNA-seq data for leaf ageing was used (Woo et al., 2016; **Supplemental Dataset S2**). This analysis revealed that *ANAC046*, *ANAC053* and *ANAC055* are most strongly induced during senescence **(Figure 3B**), suggesting that these might contribute to the increased expression of proteasomal subunit genes during leaf ageing.

**Figure 3.**
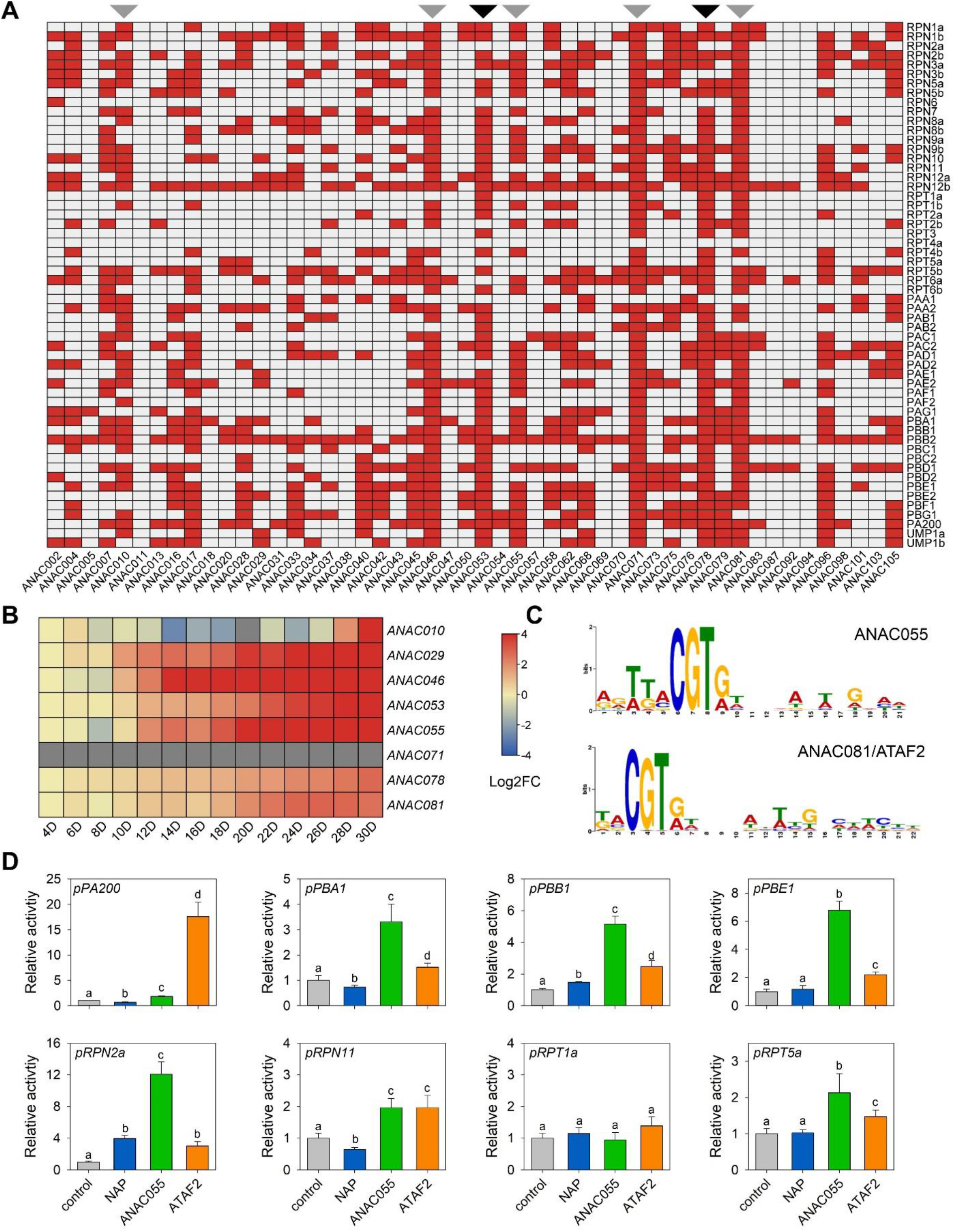
Additional NAC transcription factors can act as upstream regulators of the proteasome in Arabidopsis. A, The association of 50 NAC transcription factors with the promoter (−3000 bp) of 56 proteasomal genes was analysed by screening available DAP-seq DNA-binding data (O’Malley et al., 2016). Red tiles indicate binding, while grey indicates no binding. Black triangles above the columns indicate the known transcriptional regulators of the proteasome, ANAC053 and ANAC078. Grey triangles above the columns represent putative novel regulators of nearly all proteasomal genes. B, To reveal if the identified NAC transcription factors are induced during leaf senescence, available temporal RNA-seq data for leaf ageing was analysed (Woo et al., 2016). Heatmap represent differential expression for each timepoint as compared to the expression at day 14. C, Shown are the reported recognition motifs of ANAC055 and ANAC081/ATAF2. Data obtained from O’Malley et al. (2016). D, Transient transactivation assays with eight different proteasomal promoters controlling a luciferase reporter gene. Three NAC transcription factors were tested (ANAC029, ANAC055 and ANAC081/ATAF2). Bars represent means of four biological replicates. Significant differences were determined using one-way ANOVA with post-hoc Tukey HSD Test. Different letters indicate significant differences (p < 0.05).

Interestingly, our analysis indicates that there are potentially many other upstream regulators of the proteasome (**Supplemental Dataset S2 and S3**). Amongst the WRKY transcription factors we found that WRKY7, WRKY21, WRKY22, WRKY40, WRKY45, WRKY70 and WRKY75 interact with most proteasomal genes (**Supplemental Figure S2**). Amongst these, *WRKY45* and *WRKY75* are upregulated during leaf ageing and were shown to play a role in senescence progression (Barros et al., 2022; Guo et al., 2017). Of the 65 analysed AP2ERF transcription factors (**Supplemental Figure S3**), five (CRF10, DEAR2, ERF19, ERF34, and ERF105) associate with most proteasomal subunit genes, but only the uncharacterized *CYTOKININ RESPONSE FACTOR 10* (*CRF10*) shows an increased expression during leaf ageing. From the homeobox transcription factors is was found that BLH1, HAT1 and PDF2 recognize nearly all proteasomal subunit promoters (**Supplemental Figure S4)**, of which *BLH1* is upregulated during leaf ageing and was shown to be involved in the regulation of ABA responses (Kim et al., 2013). In addition, the HSF transcription factors HSF3 and HSFA6B are potential regulators of the proteasome, however, not during leaf ageing (**Supplemental Figure S4**). Of the 57 MYB transcription factors with reported binding data, eight (MYB31, MYB33, MYB44, MYB55, MYB67, MYB70, MYB77, and MYB107) might act as transcriptional regulators of the proteasome (**Supplemental Figure S5**). During leaf ageing both *MYB31* and *MYB33* are upregulated, which are known for their role in reproductive development and ABA responses (Shi et al., 2022; Reyes and Chua, 2007). Four senescence-induced bZIP transcription factors, ABI5, GBF3, TGA2 and TGA3, act as potential regulators of the proteasome (**Supplemental Figure S6**). TGA2 is involved in salicylic acid (SA) induced leaf senescence (Pham et al., 2022), and also TGA3 is known for its role in SA signalling (Choi et al., 2010). In contrast, ABI5 and GBF3 are involved in ABA responses and were shown to regulate leaf senescence (Su et al., 2016; Llorca et al., 2015). Furthermore, the uncharacterized and senescence-induced ZAT5 transcription factors associates with nearly all proteasomal genes (**Supplemental Figure S7**). Also, among bHLH, DOF, and Myb-like TFs upstream regulators of the proteasome were found (**Supplemental Figure S8-10**), however, none appears to be senescence-induced. Next to that, amongst TCP and G2-like transcription factors no broad proteasomal regulator was found (**Supplemental Figure S11**). Taken together, 13 senescence-induced transcription factors were found to associate with most proteasomal genes, suggesting a complex and robust regulation of the 26S proteasome during leaf ageing.

To validate the role of the here identified transcription factors in the regulation of the proteasome we focussed on NAC transcription factor members. Screening the promoters of the proteasomal genes for the NAC consensus binding sequence C[GT]TNNNNNNNA[AC]G revealed that all contain at least one (**Supplemental Figure S12**). Moreover, these corresponded also to the reported binding motifs of ANAC055 and ATAF2 (**Figure 3C**), as based on the DAP-seq analysis (O’Malley et al., 2016). To validate regulation by these NACs we employed a transient transactivation assay using the promoter of eight different proteasomal genes upstream of a luciferase reporter gene (**Figure 3D**). Next to ANAC055 and ATAF2 we also included NAP, although it does not interact with all proteasomal genes. ANAC055 and ATAF2 were able to activate the expression of seven reporters, confirming that they represent broad regulators of the 26S proteasome. NAP repressed the expression of three reporters (*pPA200*, *pPBA1* and *pRPN11*) and upregulated two (pPPB1 and pRPN1), and therefore did not act as a general promoter of proteasomal activity.

### Steady-state mRNA and ribosome-associated mRNA of proteasomal subunits during leaf senescence has no correlation to protein abundance

So far, we have shown that 26S proteasome activity and protein abundance of several subunits decreases during senescence, while at the same time the expression of proteasomal genes is induced. To understand if the decrease in subunit abundance is due to reduced translation, we compared the level of ribosome-associated mRNA with that of total mRNA in young and old leaves. To analyse the level of ribosome-associated mRNA we used TRAP technology by employing RPL18-FLAG lines (Mustroph et al., 2009). First, we compared the relative transcript change between old and young leaves at the total and polysome-bound mRNA level (**Figure 4**). Our analysis indicates that the senescence-induced expression of proteasomal subunit genes not only occurs in the total RNA pool, but also at the level of polysomes.

**Figure 4.**
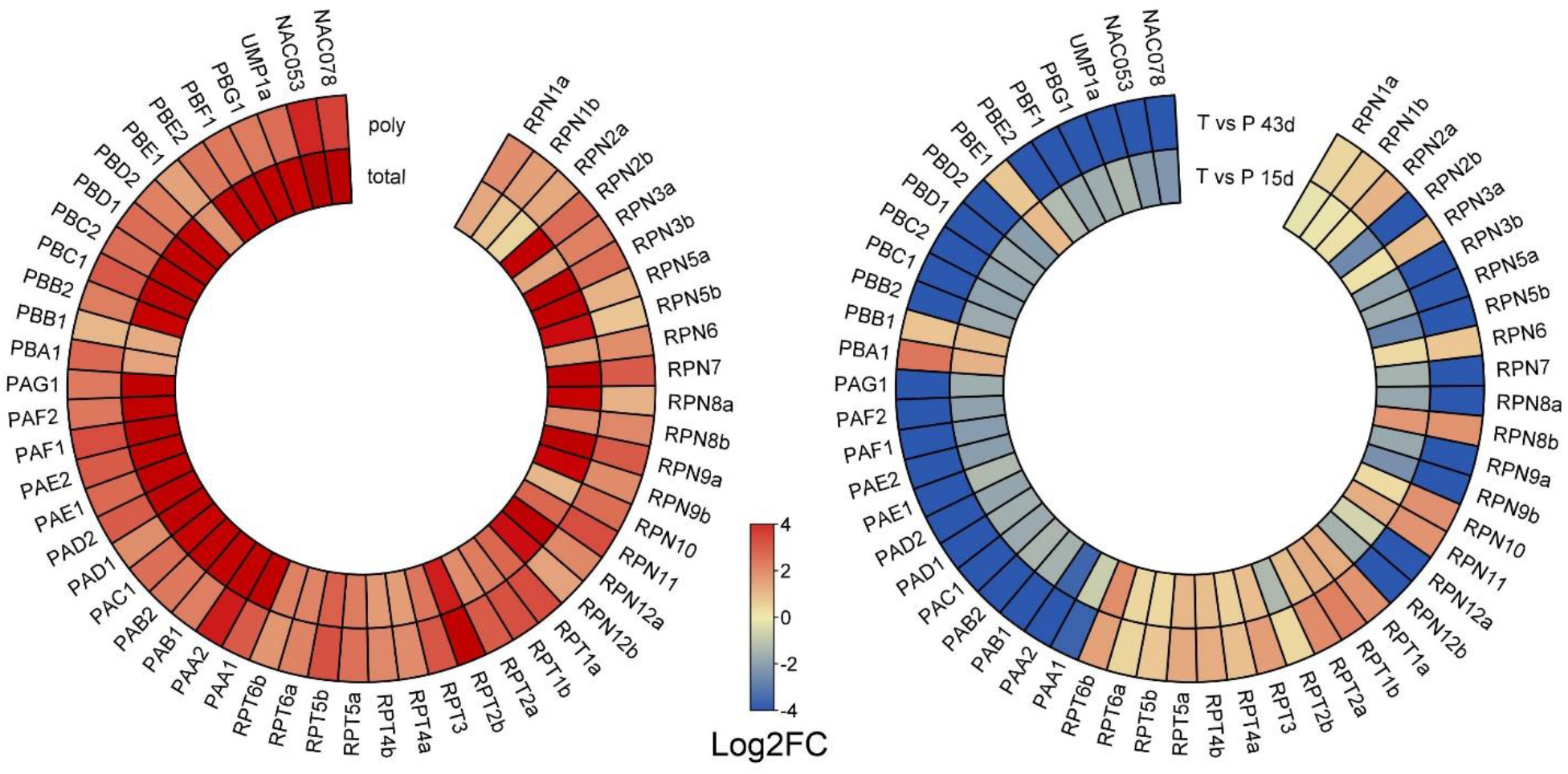
Comparison between proteasomal transcriptome and translatome during leaf ageing. The expression and polysome-bound status of mRNA for proteasomal genes at day 15 and day 43 was analysed by using Translating Ribosome Affinity Purification (TRAP). On the left, the expression of all proteasomal genes tested at the transcriptome (total) and translatome (poly) level at 43 days as compared to 15 days was plotted. On the right, the relative abundance of the mRNA at day 15 and 43 between the total RNA and the polysome-bound level was compared. Red indicates a higher relative abundance at the transcriptome level, while blue indicates an increased level at the polysome-bound level. All data presented represent the means of at least three biological replicates.

This observation suggests that in principle proteasomal subunit mRNA is still being translated during leaf senescence. Next to that, we compared the relative transcript abundance of proteasomal subunits between the total and polysome-bound mRNA pools at day 15 and day 53 (**Figure 4**). This analysis revealed differential regulation of proteasomal subunit mRNAs in the total and polysome-bound fractions. For instance, mRNAs of genes encoding for RPT subunits are more abundant at the total RNA level as compared to the polysome bound, in both young and old leaves. In contrast, mRNAs of genes encoding 20S particle subunits are more abundant in the polysome-bound fraction, as compared to the total RNA fraction, with three remarkable exceptions. The mRNAs of the proteolytic subunits of the 20S particle, PBA1, PBB1 and PBE1, are all more abundant in the total RNA fraction as compared to the polysome-bound fraction. Finally, mRNAs of RPN subunits do not show a common pattern, but some are more abundant at the polysome-bound fraction while others are more abundant in the total RNA fraction. Our data indicate that especially the RPT and proteolytic subunits are under translational control. Furthermore, the increased abundance of proteasomal mRNAs in polysomes during senescence suggest that the decreased protein abundance might be due to a faster turn-over of proteasomes during senescence.

### Transcriptional regulation of the proteasome in developing siliques and seeds

Leaf senescence results in nutrient remobilization to other parts of the plants, especially during reproductive development the developing siliques and seeds act as sinks (Guiboileau et al., 2012; Schippers et al., 2015). To understand whether the proteasome is similarly regulated in developing siliques and seeds, we assessed the expression of proteasomal genes at two different stages. Developing siliques were harvested at 10 days post anthesis (young) and 25 days (old), and prior to freezing of the material the developing seeds were collected. The expression data for siliques did not reveal a clear differential expression of proteasomal genes between the two timepoints analysed here (**Figure 5**). This although the old siliques already showed clear visual symptoms of senescence (**Supplemental Figure S13**). In contrast, during seed maturation all proteasomal genes showed a clear upregulation (**Figure 5**). Thus, there appears to be a huge difference in the transcriptional regulation of the proteasome between siliques and seeds, although the silique encloses the seed and undergoes senescence.

**Figure 5.**
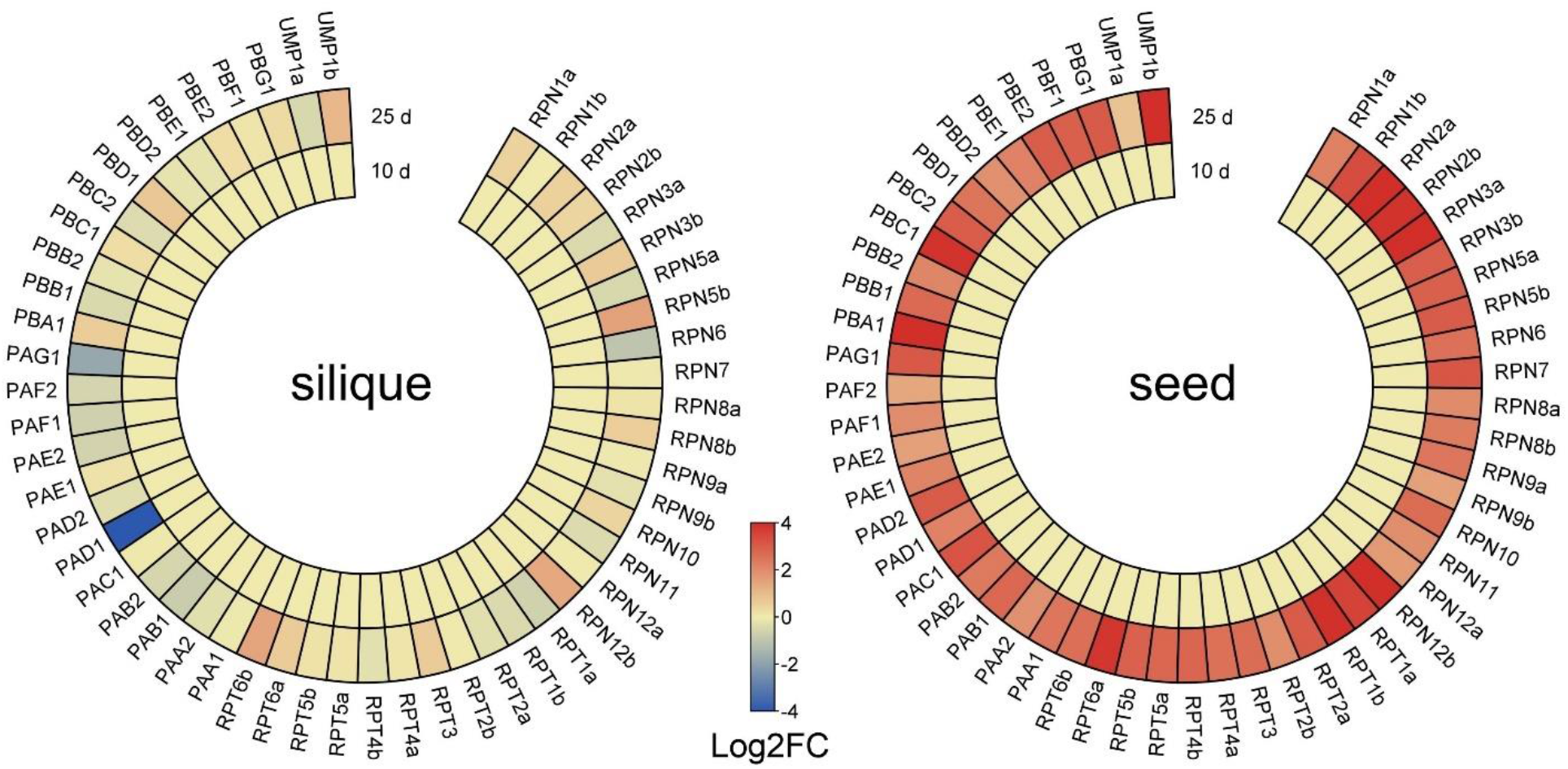
Expression of proteasomal gene in young and mature siliques and seeds. On the left, the relative expression level of the proteasomal genes in 10- and 25-day-old siliques is presented. All values were compared to the data of 10-day-old siliques. On the right, the relative expression of proteasomal genes in seeds is presented at day 10 and day 25. All values were compared to the data of 10 day-old seeds. All data presented represent the means of at least three biological replicates.

### Proteasome activity and subunit abundance in developing siliques and seeds

Recently, it was shown that during the first eight days of silique (containing seeds) formation transcript and protein abundance of several proteasomal subunits declined directly after anthesis (Yu and Hua, 2022). Here we harvested siliques and seeds separately and found no difference in expression for proteasomal genes between young and old siliques but found an increase in expression in seeds. To determine if the expression behaviour correlates with protein abundance, we first performed a western blot analysis (**Figure 6A**). Both RPN11 and PBA1 showed a decreased level in siliques and seeds of 25 days as compared to the 10-day-old samples. Thus, in both tissues there appears to be no correlation between expression and protein abundance, similar to what was found in leaves. In contrast, proteasome activity for siliques was enhanced with ageing (**Figure 6B-C**). Especially DC and SC 26S proteasome activity in ageing siliques increased while 20S activity appeared to remain unaffected. Of note, 20S activity bands of the siliques (20S +) run higher in the gel as compared to those in leaves, suggesting the presence of chaperones or other proteins interacting with the core particle in siliques (Bonea et al., 2021).

**Figure 6.**
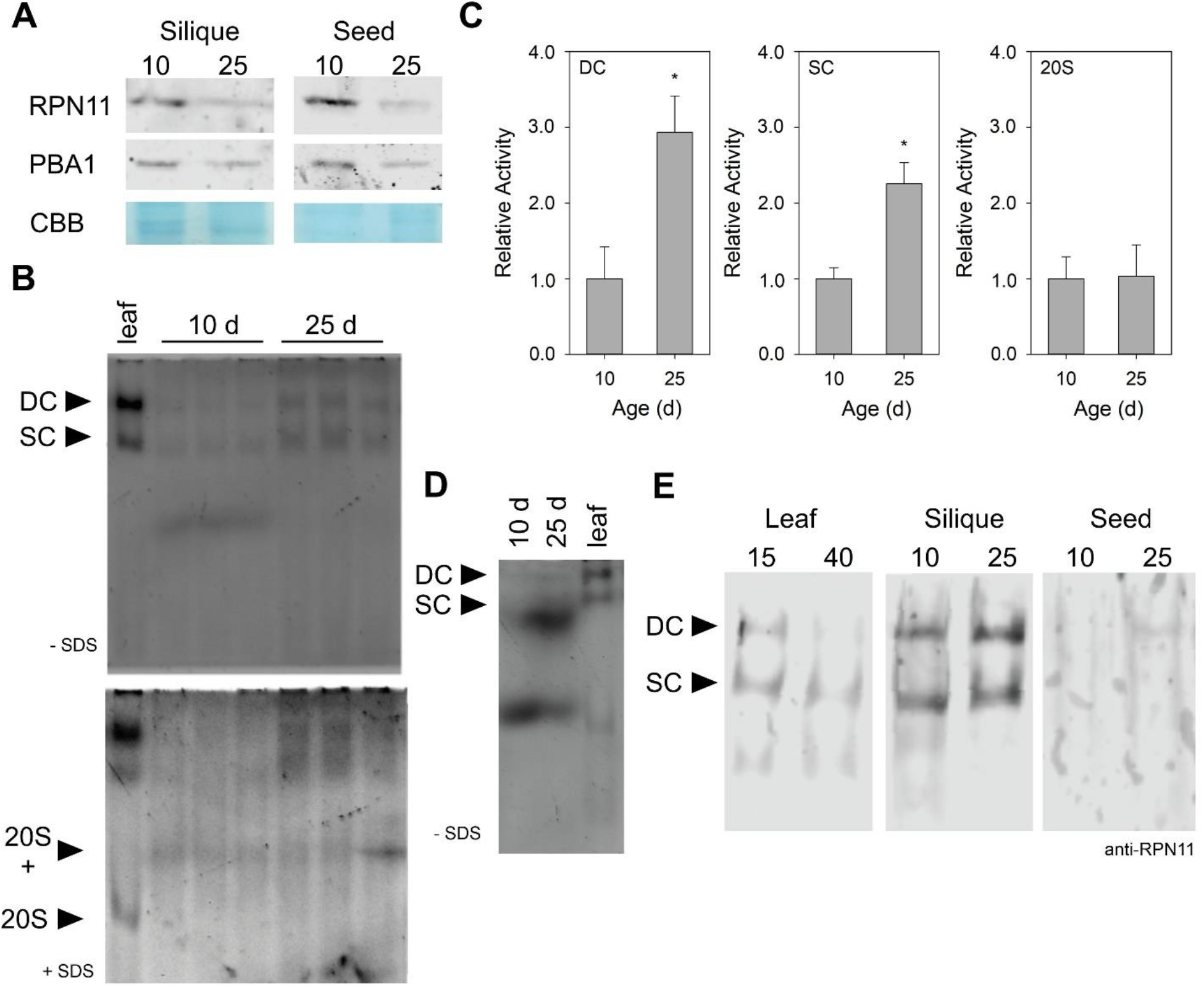
Proteasome activity and assembly increase in ageing siliques and seeds. A, Proteasomal subunit abundance in developing siliques and seeds. Total protein was separated and blotted and individual subunits were detected using specific antibodies. Coomassie Brilliant Blue (CBB) stained gels were used loading control. B, Proteasome activity assay by Suc-LLVY-AMC (100 μM) stain of total silique lysates resolved on 4% Native-PAGE (DC: double-capped 26S proteasome; SC: single-capped 26S proteasome; 20S: 20S proteasome: 20S +: 20S proteasome + putative chaperones). Total protein was isolated from at day 10 or day 25 after anthesis from siliques or seeds. Three independent biological replicates were loaded on gel. Leaf samples were taken along as a control for the staining. C, Bar graphs show relative activity for the different proteasomal complexes as based on digital quantification of band intensities from the activity gels. Data is expressed relative to the activity present at day 10, which was to 1.0. Asterisk indicates significant differences as determined by Student’s t test (p < 0.05). D, Proteasome activity assay by Suc-LLVY-AMC (100 μM) stain of total seed lysates resolved on 4% Native-PAGE. E, Immunoblot of Native-PAGE gels containing proteasomal complexes with RPN111 antibody. Total protein was isolated from leaves, siliques and seeds and the indicated days. DC: double-capped 26S proteasome; SC: single-capped 26S proteasome.

For seeds, proteasome activity gels suffered from unspecific bands (**Figure 6D**). Still, there appears to be an increase in DC and SC 26S proteasome activity for 25-day-old samples as compared to 10-day-old seeds. In order to understand why proteasome activity increases, while proteasomal subunit abundance at the protein level decreased in ageing siliques and seeds, we immunoblotted native protein activity gels with an antibody against RPN11 (**Figure 6E**). The results showed that mature siliques and seeds have increased abundance of proteasomal complexes, although mature siliques and seeds have lower RPN11 protein, indicating that proteasome assembly is most likely enhanced.

## Discussion

Previous studies have revealed that proteasome activity declines with age in animal cells, including human soma cells such as epidermal cells, fibroblasts and lymphocytes (Bulteau et al. 2000; Carrard et al. 2003; Chondrogianni et al. 2003) and rat organ cells like lung, liver, heart, kidney, spinal cord, hippocampus, cerebral cortex, muscle tissues (Bardag-Gorce et al. 1999; Bulteau et al. 2002; Conconi et al. 1996; Husom et al. 2004; Keller et al. 2000; Dasuri et al. 2009). While in germline cells, like eggs or gonads, proteasome capacity is maintained (Fredriksson et al. 2012; Tsakiri et al. 2013). Here we found that during leaf ageing 26S proteasome activity deteriorates, while in siliques and seeds an increase with age was observed. These findings are in analogy to those reported for the different animal systems, suggesting a generally conserved role for the 26S proteasome in ageing. This observation is underlined by our finding that cytokinin reverts the deterioration of 26S proteasome activity and delays senescence. In addition, we found that there is no correlation between 26S proteasome activity, proteasomal subunit expression and translation and protein level, suggesting a complex regulation of proteasomal activity *in vivo*. Finally, we report the identification of novel transcriptional regulators of the proteasome, which was demonstrated for ANAC055 and ANAC081/ATAF2, which are both induced during leaf senescence. Our results provide novel insights into the regulation of the 26S proteasome in plants, and hint at a conserved role in preventing ageing of tissues between animals and plants.

Although during leaf senescence a strong upregulation of proteasomal genes occurs at the transcriptome and translatome level, still a decrease in 26S proteasome activity and proteasomal protein abundance was observed. The increased 26S proteasome activity during the early stage of leaf development might support its crucial role in cell proliferation and differentiation (Genschik et al., 2014). As inhibition of the proteasome causes accelerated dark-induced senescence, it appears that the transcriptional and translation efforts to maintain proteasome activity during leaf senescence aims at enabling a programmed disassembly of the leaf to optimize nutrient salvage. The reduced 26S proteasome activity could be on the one hand caused by the decreased availability of ATP during senescence (Chrobok et al., 2016), since proteasome assembly and the functioning of the 26S proteasome relies on ATP (Marshall and Vierstra, 2019). On the other hand, during senescence autophagic activity increases as bulk degradation mechanism of cellular compartments (Havé et al., 2017). Plants employ the autophagy pathway to selectively degrade proteasomes through RPN10, which act as an autophagy adaptor (Marshall et al., 2015). Potentially the increased autophagic activity during leaf ageing decreases 26S proteasome abundance and thereby promotes the onset of leaf senescence. In contrast to the 26S proteasome activity, an increase in the activity of the 20S proteasome is observed during leaf senescence. Normally, the lid particle controls the selective degradation of target proteins by unfolding and presenting them to the core particle. During leaf senescence a strong increase in peptidase activity is observed, resulting in diverse short peptides that can be further processed by the 20S core particle. Furthermore, the 20S proteasome helps to remove damaged, especially oxidized proteins (Pickering et al. 2012), which might be important during the stressful last phase of the leaf. Plant cells enhance 20S proteasome capacity when 26S proteasome is impaired, like in RP mutants *rpt2a* or *rpn12a* (Kurepa et al. 2008). Moreover, an increased ratio of the 20S to 26S confer RP mutants stronger oxidative stress tolerance (Kurepa et al. 2008), implying that higher ratio of 20S to 26S maybe a protective strategy against high ROS content in senescing leaves (Woo et al. 2013).

Proteasome inhibitor treatment results in senescence and a strong transcriptional upregulation of the proteasome. It is well known that inhibition of the proteasome causes a positive transcriptional feed-back regulation to counteract the loss of proteasome activity (Gladman et al., 2016). So far, ANAC053 and ANAC078 have been identified as transcriptional regulators of the proteasome in Arabidopsis (Nguyen et al., 2013; Gladman et al., 2016). Inhibition of the proteasome was shown to induce next to NAC transcription factors, also those belonging to the WRKY and HSF transcription factor family (Gladman et al., 2016), suggesting that also other TFs might play a role in the feed-back regulation. Here we screened for additional transcriptional regulators by using previously published DAP-seq binding data (O’Malley et al., 2016). We detected 13 senescence-induced transcription factors that potentially acts as general regulators of the 26S proteasome. These include the senescence related WRKY45 and WRKY75 (Barros et al., 2022; Guo et al., 2017), TGA2 and TGA3 (Choi et al., 2010; Pham et al., 2022), and ABI5 controlling ABA responses during leaf senescence (Llorca et al., 2015). Thus, transcriptional regulation of the proteasome is more complex as previously reported, and appears to occur also in a tissue and time specific manner. Here we validated that ANAC055 and ANAC081/ATAF2 are general transcriptional activators of the proteasome, and both are senescence-induced.

Although proteasomal genes are induced at the transcriptome and translatome level during senescence, there does not appear to be a correlation with the actual proteasome activity. Also, in siliques, the transcriptional status of the proteasomal genes do not very well reflect proteasome activity or proteasomal subunit and proteasome abundance. Either the produced proteins are rapidly degraded, or the polysome-associated mRNA is stalled. Organisms have several mechanisms of co-translational mRNA surveillance (Shoemaker et al. 2012). Especially worth nothing is the fact that proteasomal genes were found to be co-translated in RNA granules in yeast (Panasenko et al., 2019). These granules were found to be smaller than and distinct from other RNA granules such as stress granules (Panasenko et al., 2019; Protter et al. 2016). Still, storage of proteasomal mRNAs in stress granules might potentially explain the recovery of proteasome activity by cytokinin. Taken together, the co-translation aspect of proteasomal mRNAs is a poorly explored concept in plants and deserves detailed attention.

Both in senescing leaves and maturating seeds, proteasomal genes are induced and their protein levels decrease. In contrast, proteasome activity appears to increase in seeds but decreases in senescing leaves. Potentially this observation is in analogy to the animal systems, in which in soma cells 26S proteasome activity drops, while in germ cells an increase is observed. In addition, protein synthesis during the initial phase of seed germination relies on long-lived mRNAs, without *de novo* transcription (Sano et al. 2015). The accumulation of mRNAs of proteasome subunit genes in mature seeds might provide the ability for protein homeostasis during the early stages of germination.

In summary, while the proteasome is implicated in many cellular processes, we provide here a role for the proteasome in the regulation of ageing. Furthermore, we identified many novel transcriptional regulators of the proteasome. In addition, a clear discrepancy between transcript levels and proteasome activity was observed, depending on the tissues and development phase studied. The molecular regulation of the 26S proteasome is highly complex and will require further dedicated studies in plants.

## MATERIALS AND METHODS

### Plant material and Growth conditions

The *Arabidopsis thaliana* (L.) accession Columbia-0 (col) was used for the transformation and wild type control. Seedlings in the plates were grown ½ strength Murashige and Skoog (MS) with 0,5% sucrose and grown under long-day conditions (16h-light/8h-dark) with a day/night temperature of 21°C/18°C. For *Arabidopsis* in soil, the plants grow in single pots under long-day with a day/night temperature of 21°C/18°C exposed to continuous white fluorescent light (120μmol m^−2^ s^−1^) during the day. For siliques and seeds in *Arabidopsis*, young (10 days after anthesis), older seeds and siliques (25 days after anthesis) were separated and collected with dissecting needle and the sample were immediately frozen in liquid nitrogen.

### Constructs and plant transformation

*pRPN11:GUS*, *pRPT1a:GUS*, *pRPT1b:GUS*, *pRPT3:GUS*, *pRPN6:GUS*, *pPA200:GUS*, *pRPT5a:GUS*, (GUS reporter) were generated by amplifying the promoter region of subunit genes with primers (supplement 4). PCR product was subcloned into pENTR-D for sequence validation and then ligated into pKGWFS7 (Karimi et al. 2002). The recombination constructs were transformed into Col-0 using the floral-dip method (Logemann et al. 2006).

### Histochemical Staining

GUS staining was performed as described by Jefferson et al (1987). Briefly, for GUS staining, transgenic *Arabidopsis* were immersed in GUS staining buffer (solution: 0.1M sodium phosphate buffer (pH 7.0), 10mM EDTA, 0.1% Triton X-100, 1.0mM K_3_ Fe (CN)_6_, 2mM 5-bromo-4-chloro-3-indolyl-*β*-D-glucuronide (X-Gluc) and vacuumized for 5min. After incubation overnight at 37°C, 70% ethanol was used for extraction of chlorophyll of stained plant tissues.

### Proteasome inhibitor and cytokinin treatment

Detached leaves from first pair of 29 days-old wild type Arabidopsis leaves were floated in the solution (water+DMSO), 30μM MG132 (N-(benzyloxycarbonyl)-leucinyl-leucinyl-leucinal; SelleckChem), 10μM Cytokinin+DMSO, 30μM MG132 + 10μM Cytokinin solution respectively, and incubated under darkness until leaves turned yellow, followed by harvesting samples for the subsequent experiments. For examination of GUS staining patterns under proteotoxic stress caused by MG132, surface sterilized seeds were stratified in the dark at 4°C for 2 d, and germinated on plates with or without 15μM MG132. For proteasome activity profiles under proteasome inhibitor treatment, 10 days old wild type seedlings were transferred to ½ MS medium with/without 30μM of MG132, 2μM Bortezomib for two days. The seedlings in plates were grown under long-day conditions (16h-light/8h-dark) with a day/night temperature of 21°C/18°C.

For cytokinin treatment, 30 days-old wild type *Arabidopsis* leaves were sprayed with water and 80 μM 6-BA one time per day for one week. First and second pair of 36days-old leaf were harvested for proteasome activity analysis. 11 days old wild type *Arabidopsis* seedlings were transferred to ½ MS medium containing 4μM cytokinin and grown for 4 days, shoots were harvested for proteasome activity analysis.

### Chlorophyll content analysis

According to the method (Bresson et al 2018), 10mg leaves were collected from the first pair leaves and chlorophyll were extracted by incubation in 1ml of 80% acetone overnight in darkness. The chlorophyll levels were detected by spectrophotometric measurements with absorbance at 645nm and 663nm. Chlorophyll concentration was calculated via formulas (C_T_=20.2*D645+8.02*D663) and then normalized to fresh weight (mg Chl g^−1^ FW).

### LLVY-AMC in-gel activity assay, In-solution peptidase activity assay and immune-blotting of Native Gels

Natvie gel preparation and needed solutions were made according to method with modification (Roelofs et al. 2018; Levin et al. 2018). Protein extracts were prepared according to method with small modifications (Üstün et al. 2013). Briefly, Samples were ground with liquid nitrogen and then dissolved in 100 μL fresh extraction buffer (50 mM HEPES-KOH, pH 7.2, 2 mM ATP, 2 mM dithiothreitol, and 250 mM Suc). After native gel electrophoresis, carefully dislodged the native gel into solution (20mL solution: 1.0 ml Tris-HCl (pH7.5) with 100 μl 1M MgCl_2_, 40 μl 0.5 M ATP and 100 μl 10 mM suc-LLVY-AMC), incubated gel for 30 minutes at 30°C and then used UV transilluminator for analysis. For the seed and silique proteasome capacity measurement, 5μg and 50μg total protein were used. For immune-blotting, proteins in native gel were transferred to Nitrocellulose membranes, 26S proteasome were detected with anti-RPN11.

### RNA Extraction and genes expression Analysis

Total RNA was extracted from first pair leaf from *Arabidopsis* grown in long day conditions. TRI-reagent (Sigma-Aldrich) was used for RNA extraction according to manufacturer’s protocols. Subsequently, total RNA was treated with DNase to remove genomic DNA by using TURBO DNA-free Kit (Ambion). Two microgramm of total RNA was reverse-transcribed by the Revert Aid First Strand cDNA Synthesis Kit (Thermo Fisher Scientific). Powerup SYBR Green Master Mix (Applied Biosytems) was used to commence qRT-PCR. Gene specific primers and Actin2 primers (reference gene) sequence can be found in the supplement 4. Relative expression levels for each target genes were calculated via the comparative CT method (Schmittgen et al. 2008).

### Protein extraction and western blot analyses

Crude extracts isolated with extraction buffer (150mM NaCL, 50mM Tris pH 8.0, 1% Triton X100, protease inhibitor,) were used for western blot analysis with Laemmli sample buffer (LAEMMLI, 1970). Protein samples were separated through 7.5% SDS-polyacrylamide gels under denaturing conditions and then were transferred onto Protran nitrocellulose membrane (Whatman, Kent, UK). Before the transfer, the membrane was blocked for 1h with 5% blocking buffer and incubated for 1h with primary rabbit polyclonal antibody (Afrisera, Sweden) directed against 26s proteasomal subunits. The membranes were washed twice in TBST (TBS containing 0.1% Tween-20) for 10 min each, followed by 10 min washing with 5% blocking solution, then incubated for 1h with secondary antibody against the primary antibody (Agrisera, Sweden) at room temperature. Antibodies were diluted in 5% blocking buffer according to instructions from manufacturers. Signal intensities were detected via streptavidin horseradish peroxidase (HRP) and exposure to negative films.

### Transaction assay/Dual-Luciferase Assays

The dual-Luciferase assay was used to investigate the potential regulator of 26s proteasome. After ligation into pENTR-D (Invitrogen) and sequence validation, promoter sequences were recombined into the vector pGWL7.0, while the CDS was transferred into the p2GW7 vector (Karimi et al. 2002). Protoplasts were isolated from Wild type Arabidopsis plants according to Tape-Arabidopsis Sandwich method (Wu et al. 2009). Transformation and measurement of luciferase signal with the Dual-luciferase® Reporter Assay System (Promega) were done as reported previously (Nguyen et al. 2013). For each action, four replicates were performed.

### Immuno-purification of Ribosomes

Ribosomes were isolated from *Arabidopsis* leaves according to Mustroph et al. (2009). Briefly, frozen leaf tissue was homogenized with 2mL freshly prepared polysome extraction buffer PEB (200 mM Tris-HCl, pH 9.0, 200 mM KCl, 25 mM EGTA, 35 mM MgCl_2_, 1mM PMSF, 5mM dithiothreitol (DTT), 50 μg mL^−1^ cycloheximide, 50 μg mL^−1^ chloramphenicol, 1% Detergent mix (1% (v/v)Triton X-100, 1% (v/v) Tween 20, 1% (w/v) Brij-35, 1% (v/v) Igepal CA-630), 0.5 mg mL^−1^ heparin, 1% Poly-oxyethylene 10 tridecyl ether (PTE), 1% Sodium deoxycholate (DOC)). Homogenates were clarified by centrifugation at 16000g for 10min and 10% of the clarified extract was saved for total RNA isolation, and approximately 250-300 OD600 units of the supernatant were incubated with 50 μL anti-FLAG affinity gel (Sigma) at 4℃ for 2h with gentle shaking. The affinity gel was firstly washed with PEB buffer and washed with wash buffer (200 mM Tris-HCl, pH 9.0, 200 mM KCl, 25 mM EGTA, 35 mM MgCl_2_, 1mM PMSF, 5mM dithiothreitol (DTT), 50 μg mL^−1^ cycloheximide, 50 μg mL^−1^ chloramphenicol, 20u mL^−1^ RNAsin) for additional three times. To elute the affinity-purified ribosomes, add to the beads 300 μL of wash buffer containing 200 ng μL^−1^ of FLAG_3_ peptide and 20u mL^−1^ RNAsin and incubate 30 min at 4℃ with shaking.

### Affinity purification of proteasome

The complementary transgenic line (PAG1-FLAG/*pag1-1*) were used for the affinity purification of the *Arabidopsis* proteasome (Book et al. 2010; Marshall et al. 2017). Briefly, approximately 0.5 g leaf were ground in liquid nitrogen and extracted with buffer A [50 mM 4-(2-hydroxyethyl)-1-piperazineethanesulfonic acid (HEPES)-KOH, pH 7.5, 50 mM NaCl, 10% glycerol, 10 mM MgCl_2_, 5 mM dithiothreitol (DTT), 2 mM phenylmethylsulfonyl fluoride (PMSF), protease inhibitor cocktail (P9599; Sigma) with 20 mM ATP for 20 min. The lysate was clarified by centrifugation at 18000 x g for 30 min at 4 °C and then incubated with pre-equilibrated anti-FLAG® M2 resin (Sigma; M8823). The suspension was incubated for 1h at 4 °C on a rotary shaker. After washing four times using buffer A with 20mM ATP, bound proteins were eluted using buffer A containing 500ng/μL 3X FLAG^®^ peptide (Sigma; F4799) for followed SDS-PAGE.

## Acknowledgements

The authors wish to thank the China Scholarship Council for their support. We thank Dr Richard S Marshall and Dr Richard D. Vierstra (Washington University in St Louis, USA) for providing the *PAG1-FLAG pag1-1* seeds. This project was financially supported by grants from

## Author contributions

H-W and J.H.S designed research; H-W performed research; Joost van Dongen supervise the project, gave suggestions and reviewed the manuscript; H-W and J.H.M.S wrote the paper.

