## Supplemental Figures for "Proteasomal activity is differentially regulated in source and sink tissues of Arabidopsis"

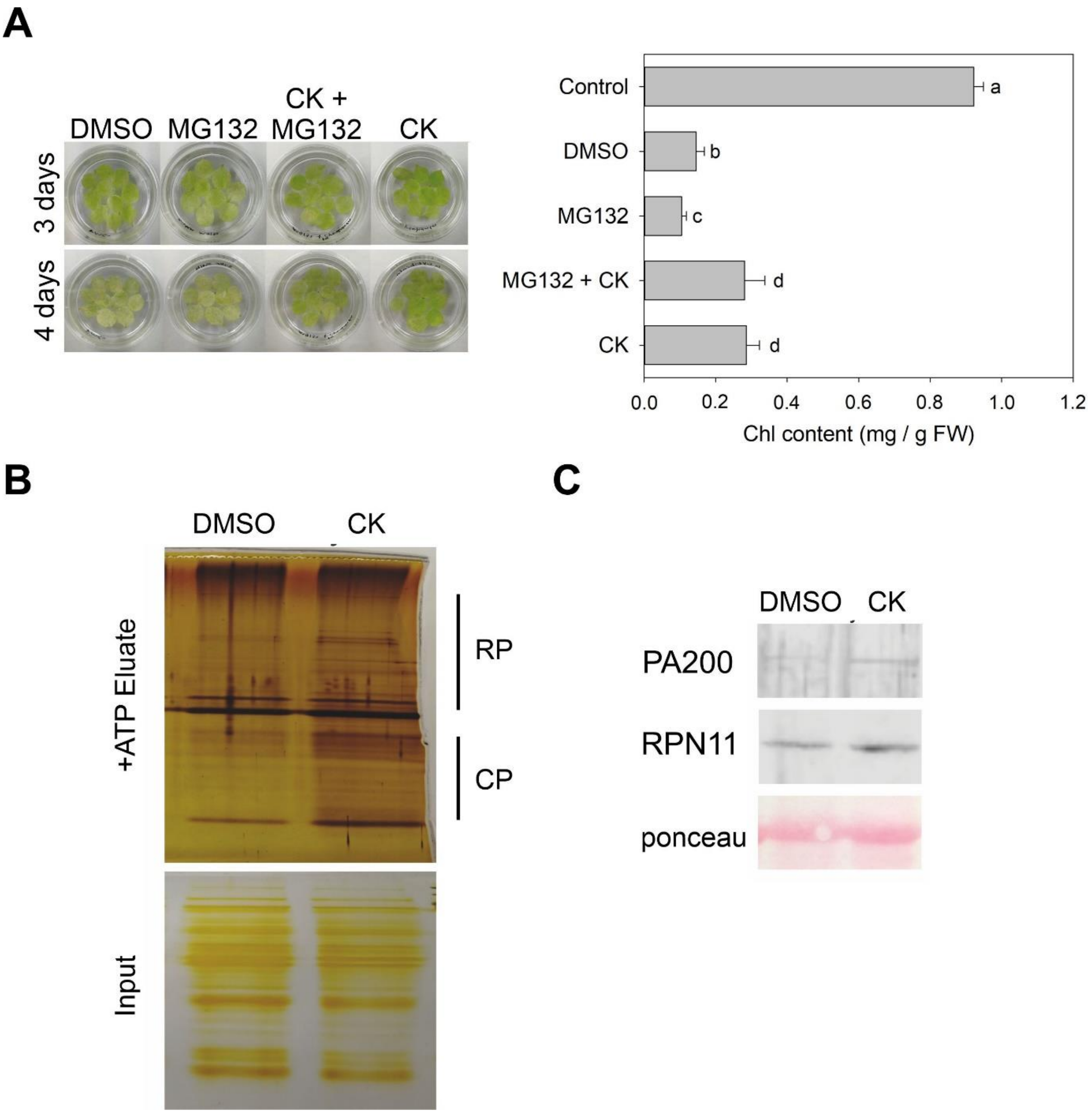


**Supplemental Figure S1. Cytokinin promotes proteasomal subunit abundance and delays senescence.** A, Dark-induced leaf senescence was performed with first leaves isolated from 30-d-old Arabidopsis plants. Leaves were floated on water containing DMSO, 30µM MG132, 10 µM CK or 30µM MG132 + 10 µM CK. Pictures show leaves after 3 and 4 days of incubation in the dark. Chlorophyll content was determined as indicated in the panel on the right. Bars represent chlorophyll content and different letters indicate significant differences according to one-way ANOVA and post-hoc Tukey HSD Test ($p < 0.05$). B, Affinity purification of proteasomes from PAG1-FLAG *pag1-1* plants treated with DMSO or cytokinin (CK). The input protein, as well as the eluate in the presence of ATP was analysed on an SDS-PAGE gel. The protein gels were stained with silver. The location of the RP and CP subunits is indicated on the right. C, proteasomal subunit abundance after cytokinin treatment. The abundance of selected subunits was detected using specific antibodies. ponceau stained blots were used loading control.

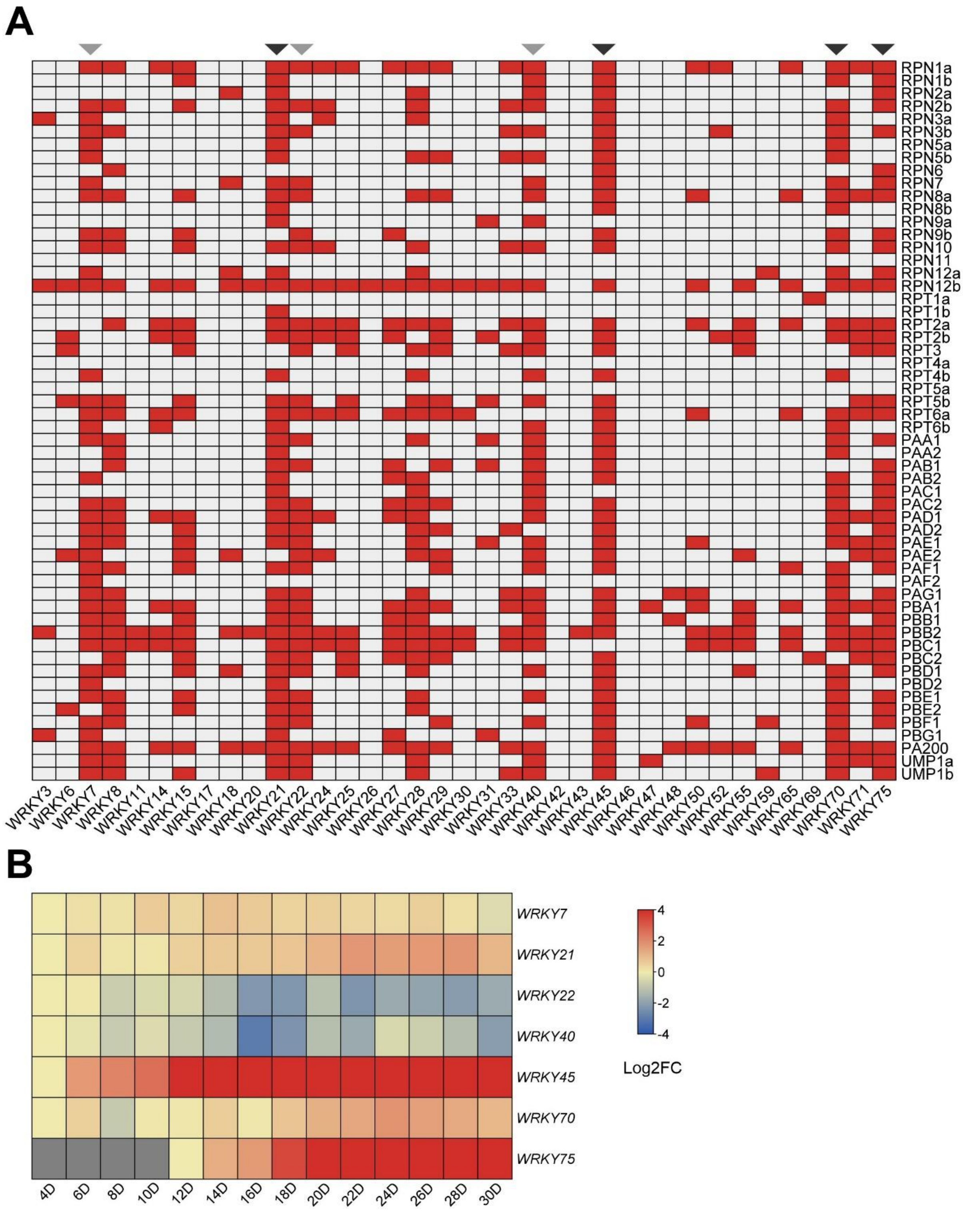


**Supplemental Figure S2. WRKY TFs involved in the transcriptional regulation of the 26S proteasome.** A, Heatmap indicating association of WRKY TFs with the promoter of proteasomal genes. The indicated binding events (red color) are based on the binding data reported by O'Malley et al. (2016). B, Heatmap indicating the expression of putative proteasome-regulating WRKY TFs during leaf ageing (Woo et al., 2016).

**A**

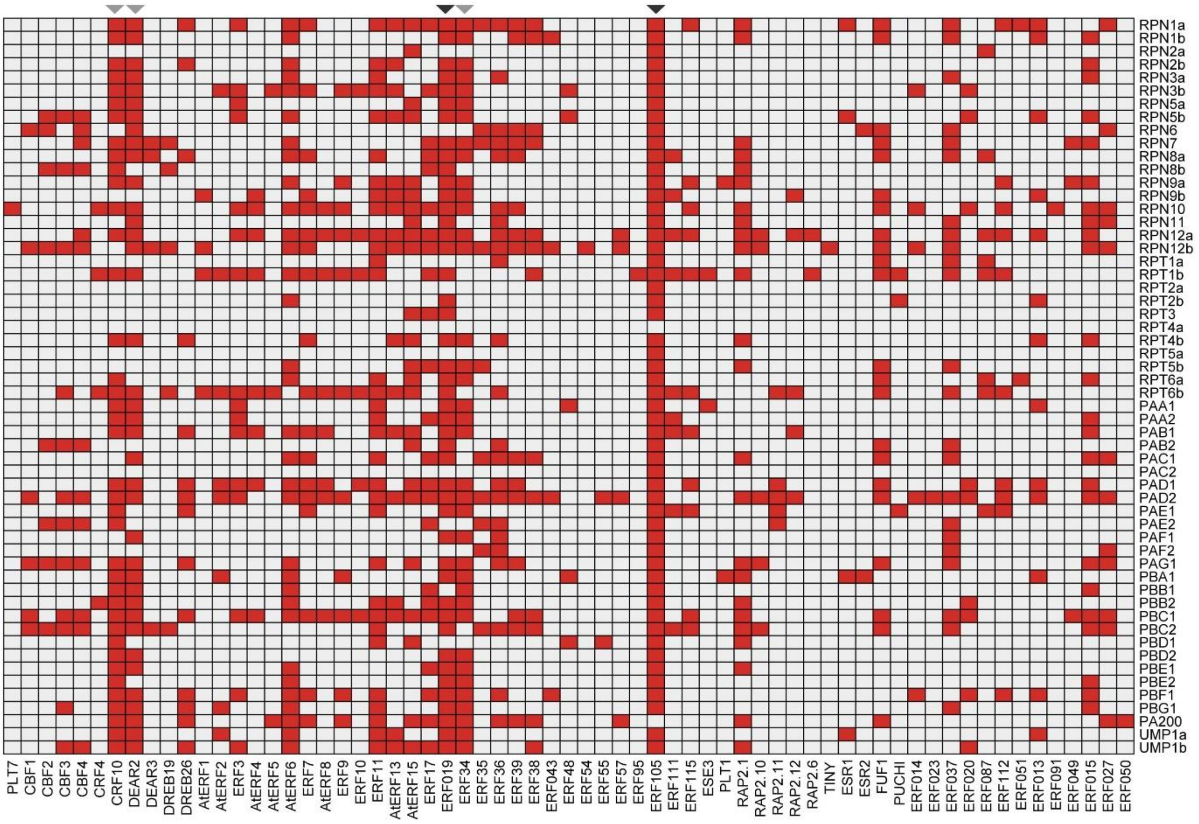

**B**

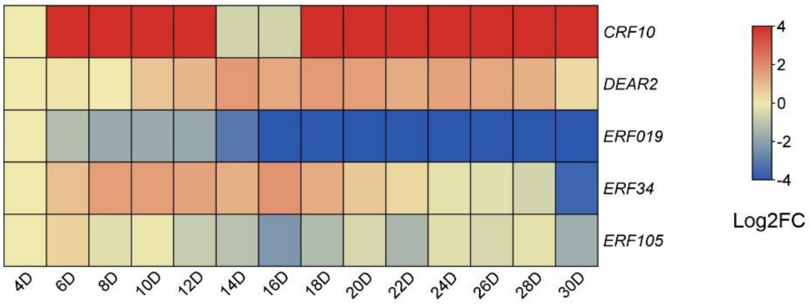


**Supplemental Figure S3. AP2ERF TFs and transcriptional regulation of the 26S proteasome.** A) Heatmap indicating association of WRKY TFs with the promoter of proteasomal genes. The indicated binding events (red color) are based on the binding data reported by O'Malley et al. (2016). B) Heatmap indicating the expression of putative proteasome-regulating WRKY TFs during leaf ageing. (Woo et al., 2016).

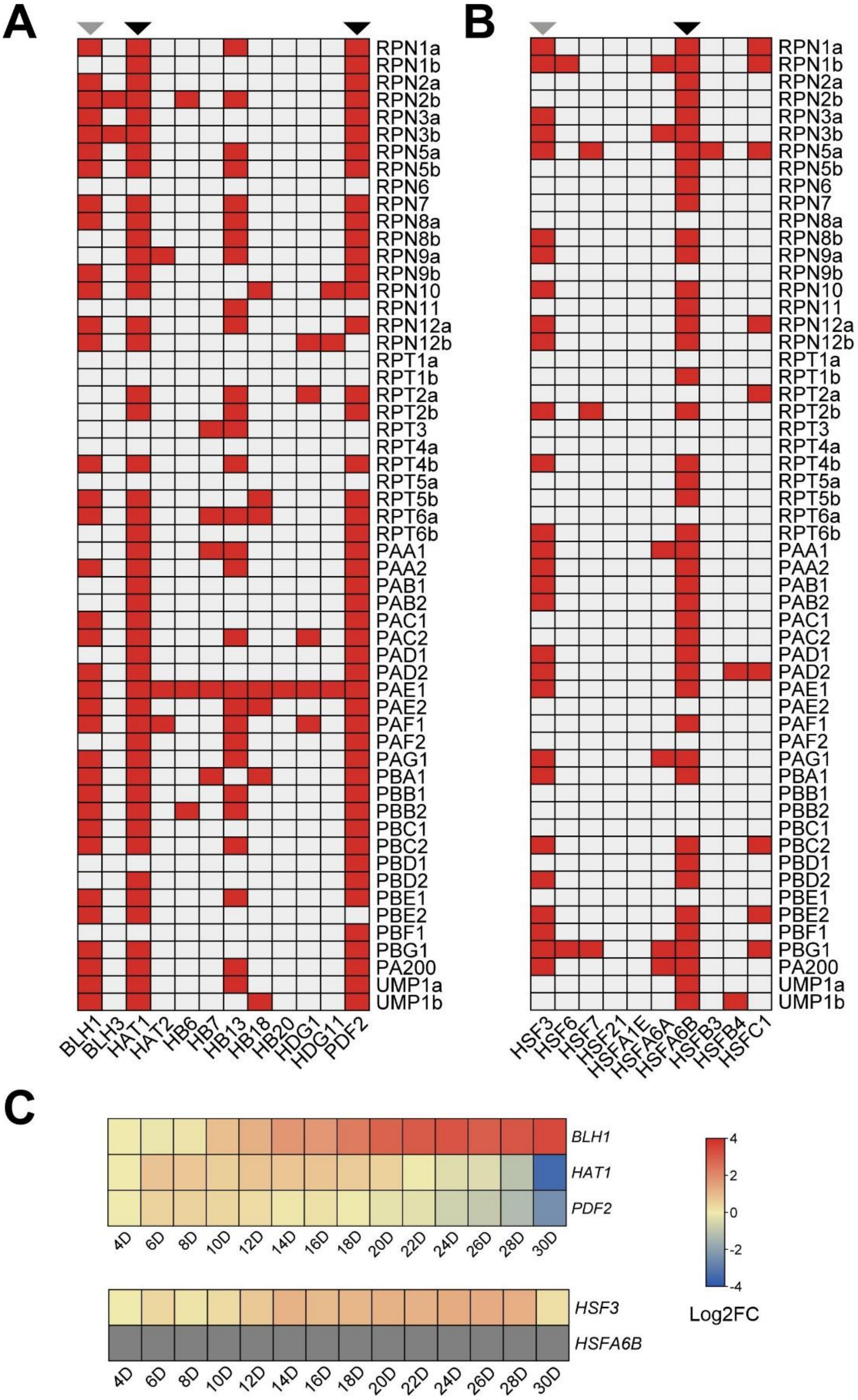


**Supplemental Figure S4. Overview of Homeobox and HSF TFs potentially involved in the transcriptional regulation of the 26S proteasome.** Heatmap indicates association of A) Homeobox and B) HSF TFs with the promoter of proteasomal genes. The indicated binding events (red color) are based on the binding data reported by O'Malley et al. (2016). C) Heatmap indicating the expression of putative proteasome-regulating TFs during leaf ageing. (Woo et al., 2016).

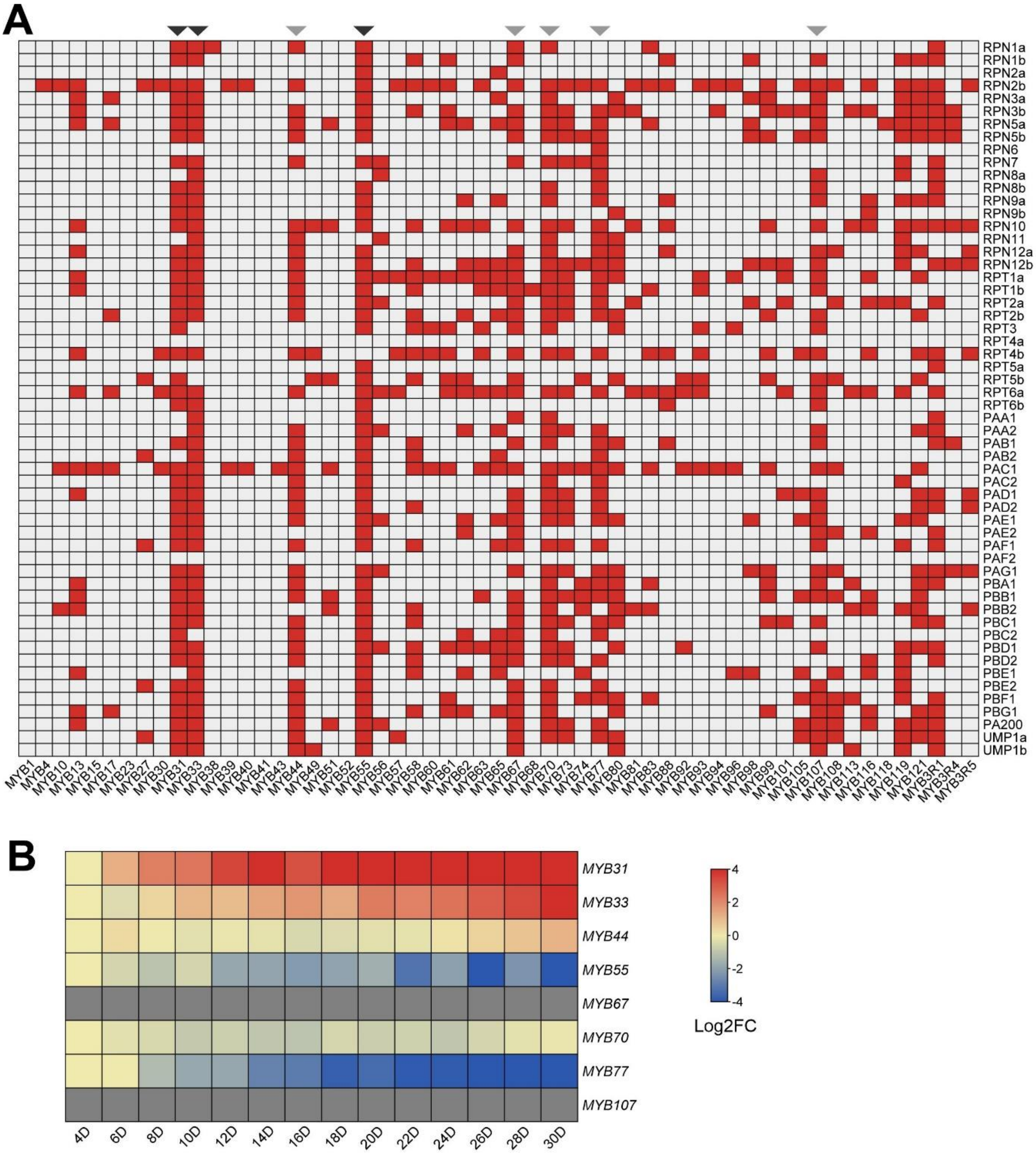


**Supplemental Figure S5. Overview of MYB TFs potentially involved I the transcriptional regulation of the 26S proteasome.** A, Heatmap indicating association of MYB TFs with the promoter of proteasomal genes. The indicated binding events (red color) are based on the binding data reported by O'Malley et al. (2016). B, Heatmap indicating the expression of putative proteasome-regulating MYB TFs during leaf ageing. (Woo et al., 2016).

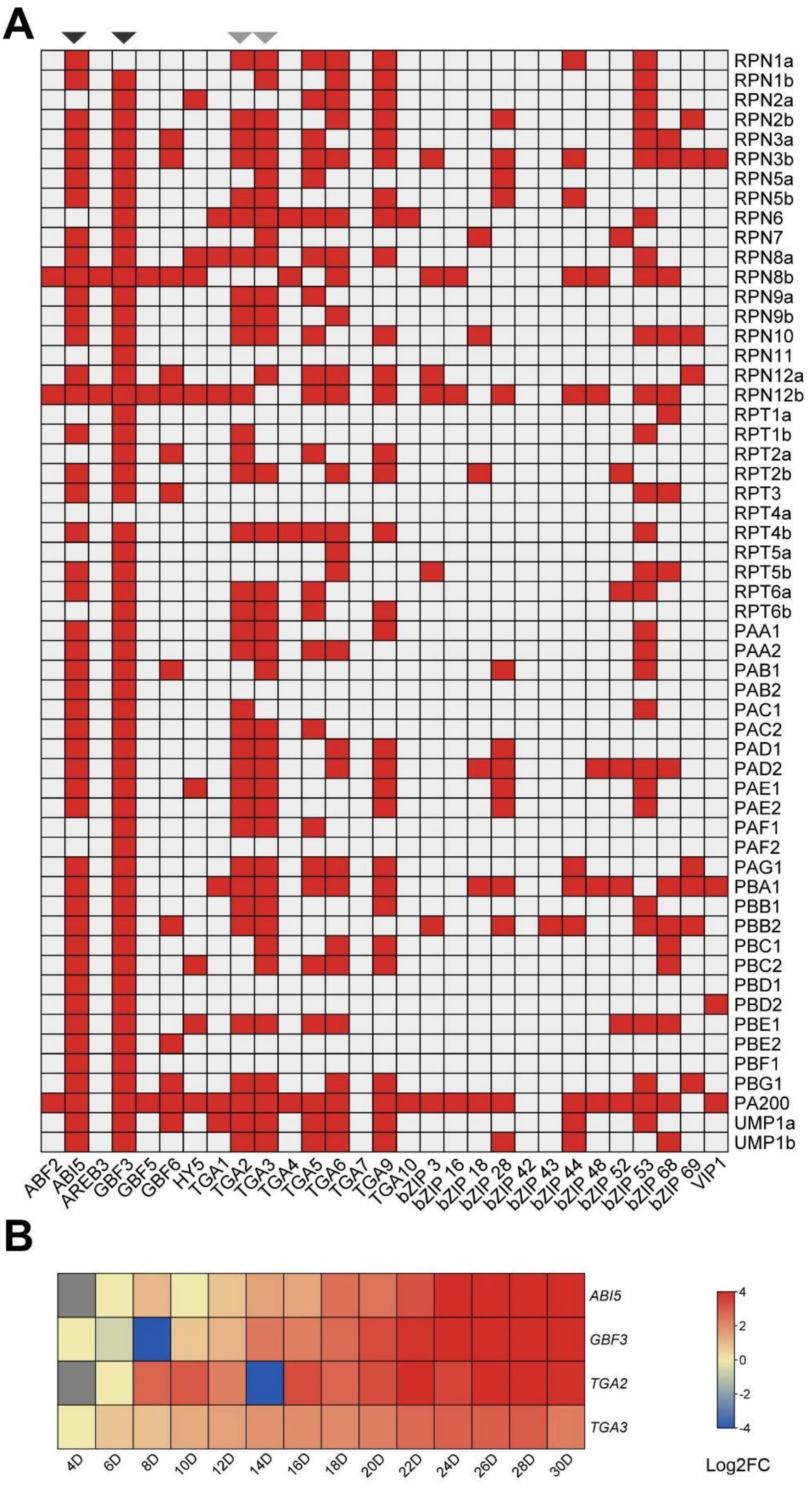


**Supplemental Figure S6. Identification of bZIP TFs involved in the transcriptional regulation of the 26S proteasome.** A, Heatmap indicating association of bZIP TFs with the promoter of proteasomal genes. The indicated binding events (red color) are based on the binding data reported by O'Malley et al. (2016). B, Heatmap indicating the expression of putative proteasome-regulating bZIP TFs during leaf ageing. (Woo et al., 2016).

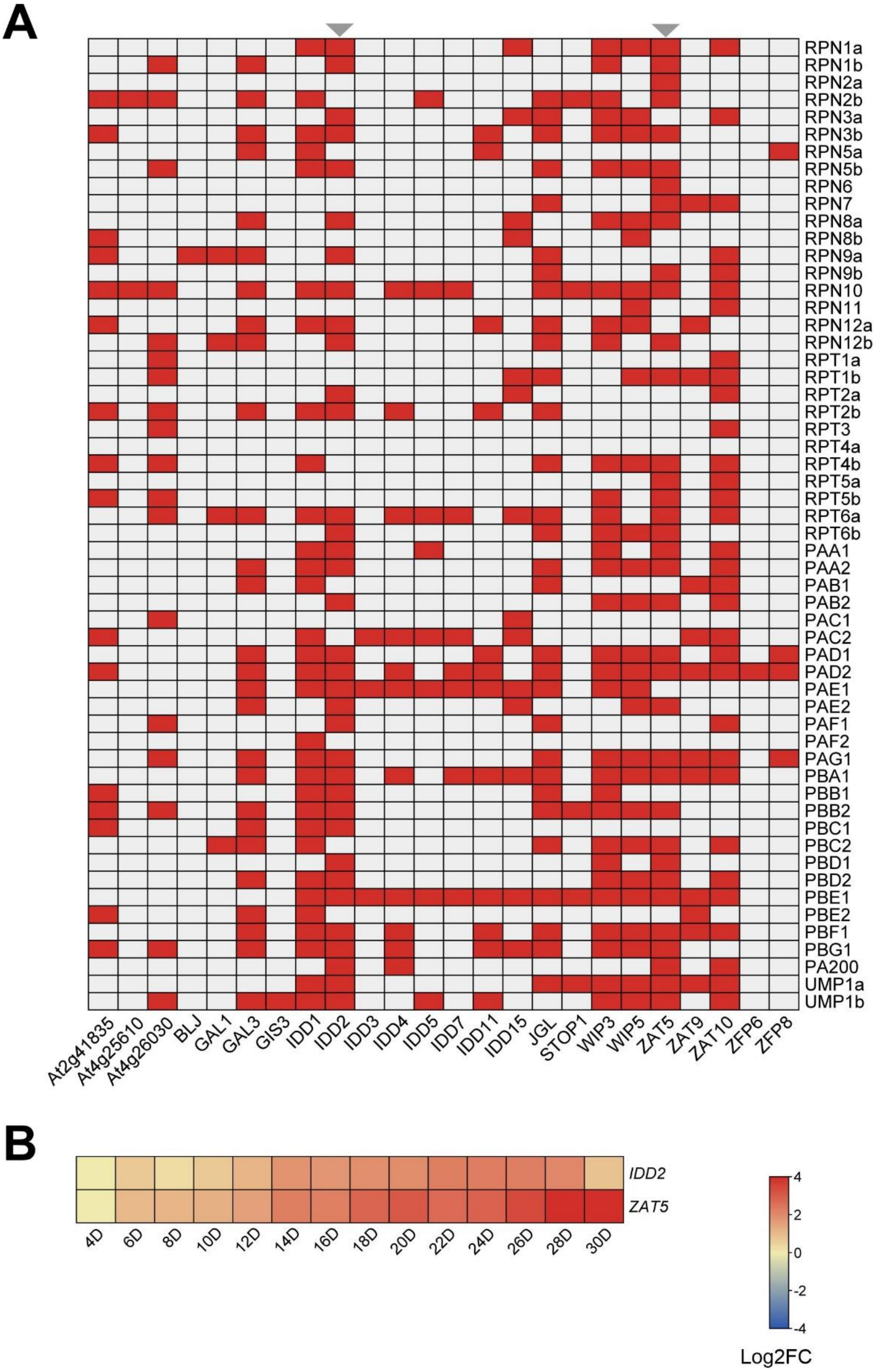


**Supplemental Figure S7. Overview of C2H2 TFs potentially involved I the transcriptional regulation of the 26S proteasome.** A) Heatmap indicating association of C2H2 TFs with the promoter of proteasomal genes. The indicated binding events (red color) are based on the binding data reported by O'Malley et al. (2016). B) Heatmap indicating the expression of putative proteasome-regulating C2H2 TFs during leaf ageing. (Woo et al., 2016).

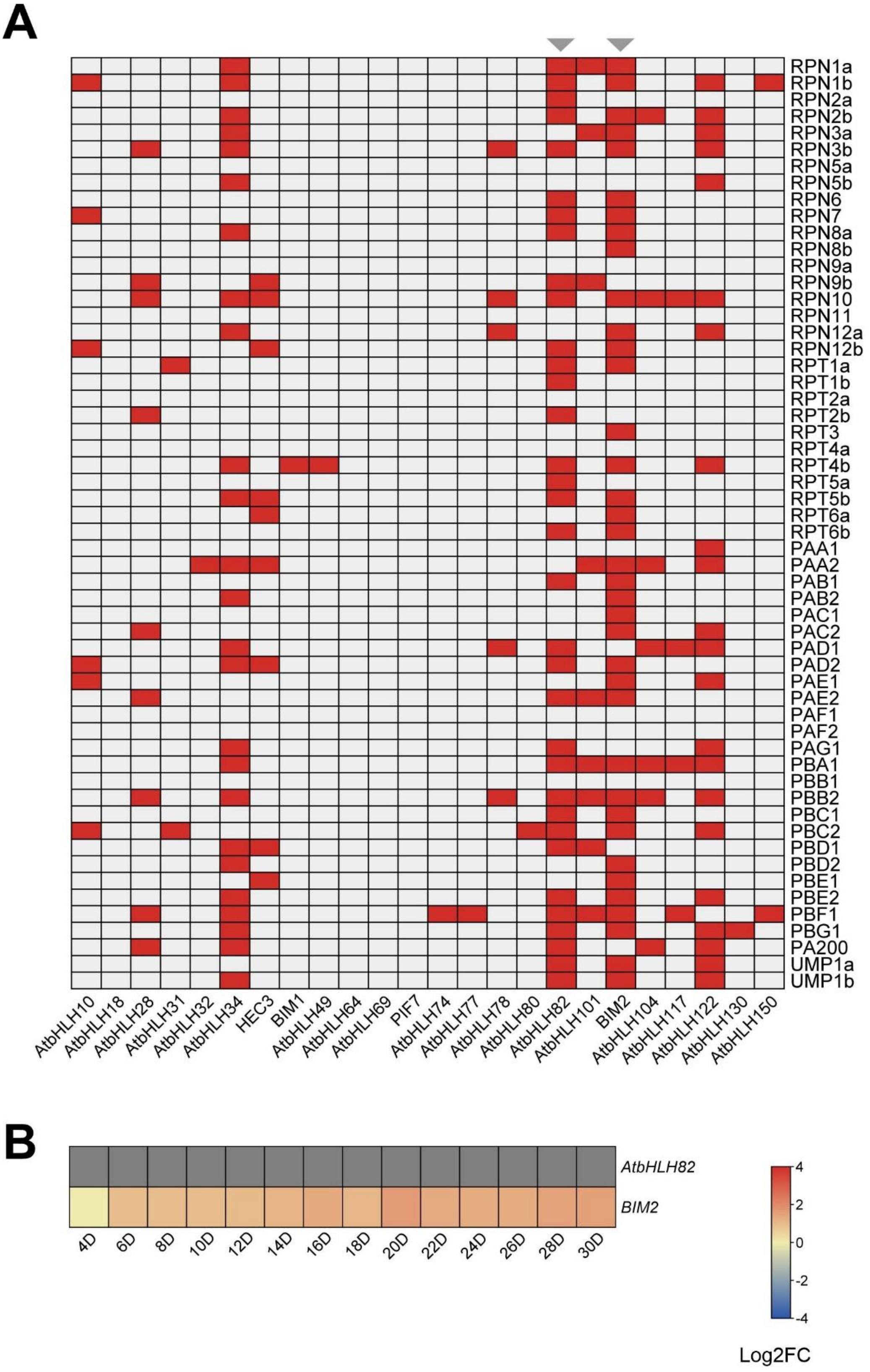


**Supplemental Figure S8. Overview of bHLH TFs potentially involved I the transcriptional regulation of the 26S proteasome.** A) Heatmap indicating association of bHLH TFs with the promoter of proteasomal genes. The indicated binding events (red color) are based on the binding data reported by O'Malley et al. (2016). B) Heatmap indicating the expression of putative proteasome-regulating bHLH TFs during leaf ageing. (Woo et al., 2016).

A

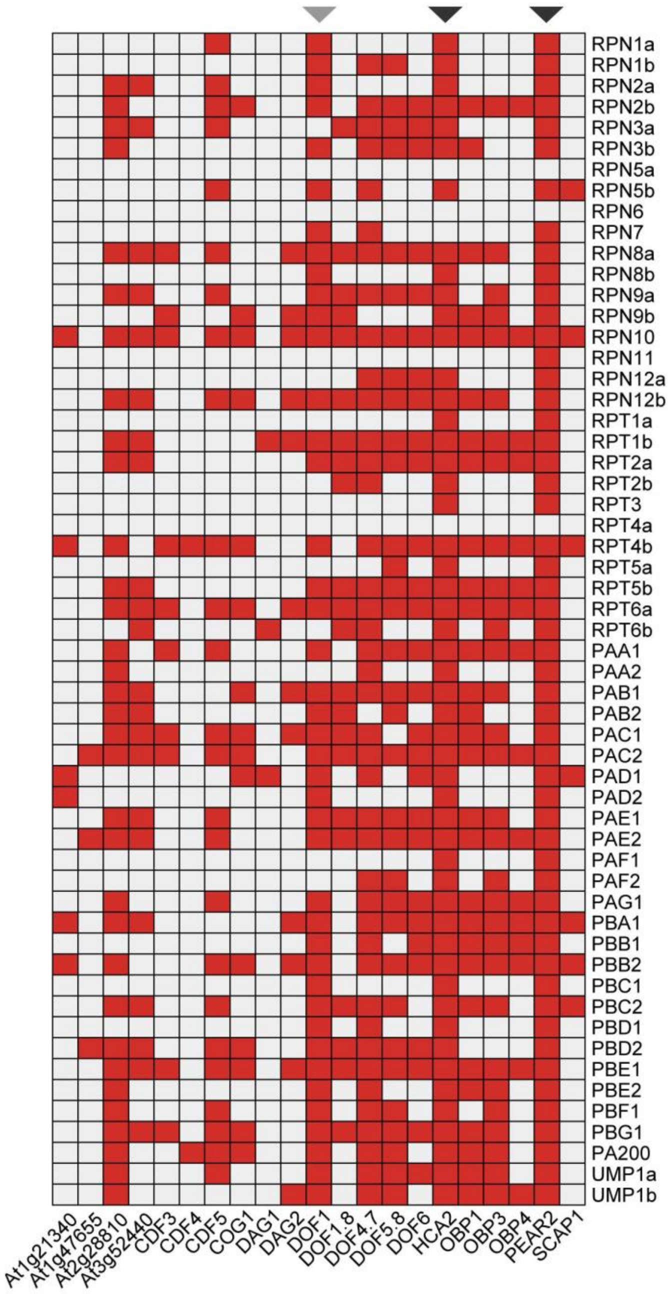


B

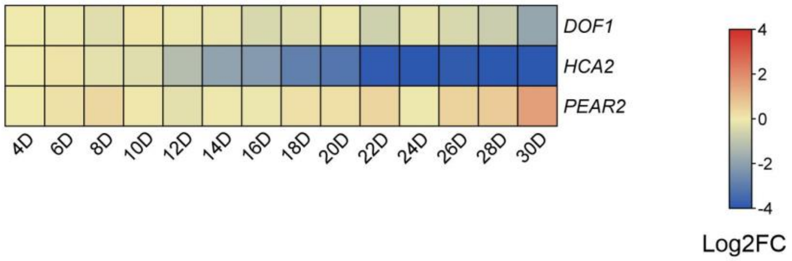


**Supplemental Figure S9. Overview of DOF TFs potentially involved I the transcriptional regulation of the 26S proteasome.** A) Heatmap indicating association of DOF TFs with the promoter of proteasomal genes. The indicated binding events (red color) are based on the binding data reported by O'Malley et al. (2016). B) Heatmap indicating the expression of putative proteasome-regulating DOF TFs during leaf ageing. (Woo et al., 2016).

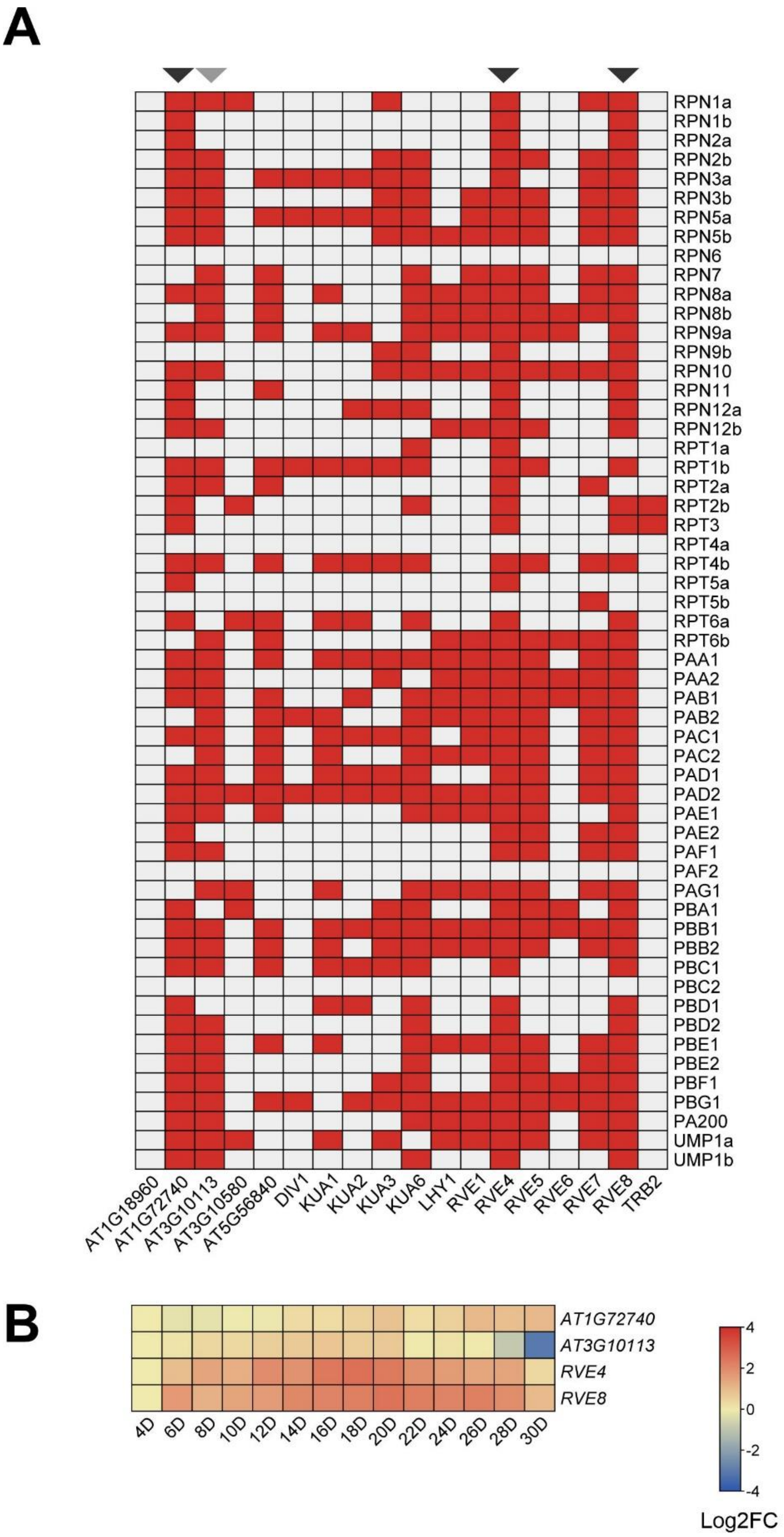


**Supplemental Figure S10. Overview of Myb-like TFs potentially involved I the transcriptional regulation of the 26S proteasome.** A) Heatmap indicating association of Myb-like TFs with the promoter of proteasomal genes. The indicated binding events (red color) are based on the binding data reported by O'Malley et al. (2016). B) Heatmap indicating the expression of putative proteasome-regulating Myb-like TFs during leaf ageing. (Woo et al., 2016).

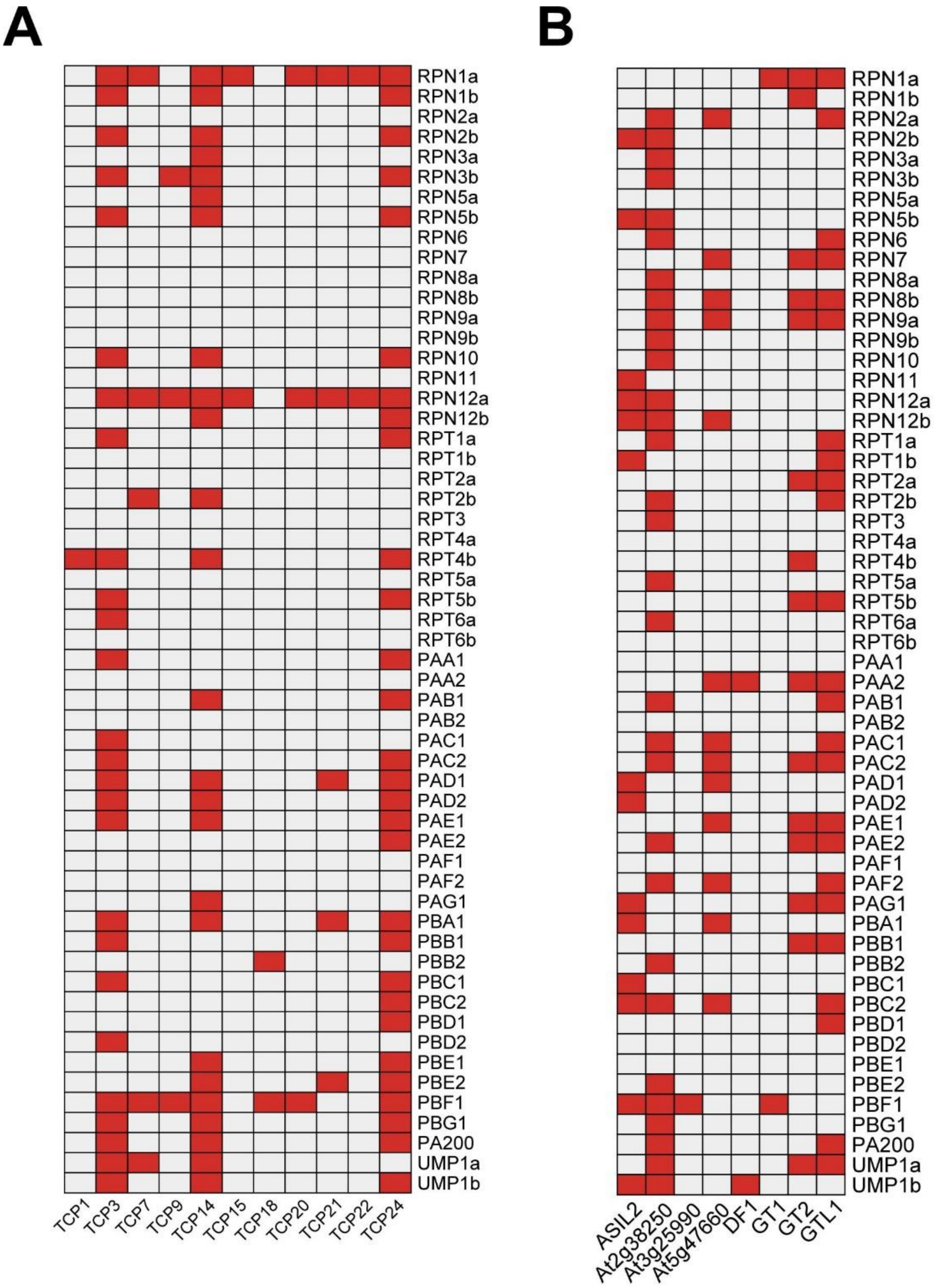


**Supplemental Figure S11. Overview of TCP and Trihelix TFs potentially involved I the transcriptional regulation of the 26S proteasome.** A) Heatmap indicating association of WRKY TFs with the promoter of proteasomal genes. The indicated binding events (red color) are based on the binding data reported by O'Malley et al. (2016). B) Heatmap indicating the expression of putative proteasome-regulating WRKY TFs during leaf ageing. (Woo et al., 2016).

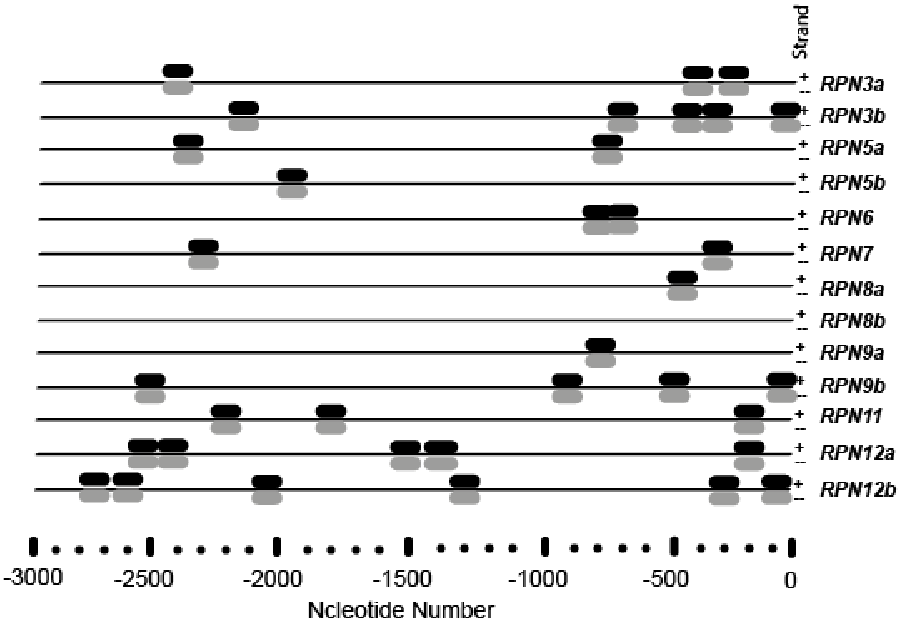
Strand
RPN3a
RPN3b
RPN5a
RPN5b
RPN6
RPN7
RPN8a
RPN8b
RPN9a
RPN9b
RPN11
RPN12a
RPN12b
-3000
-2500
-2000
-1500
-1000
-500
0
Ncleotide Number

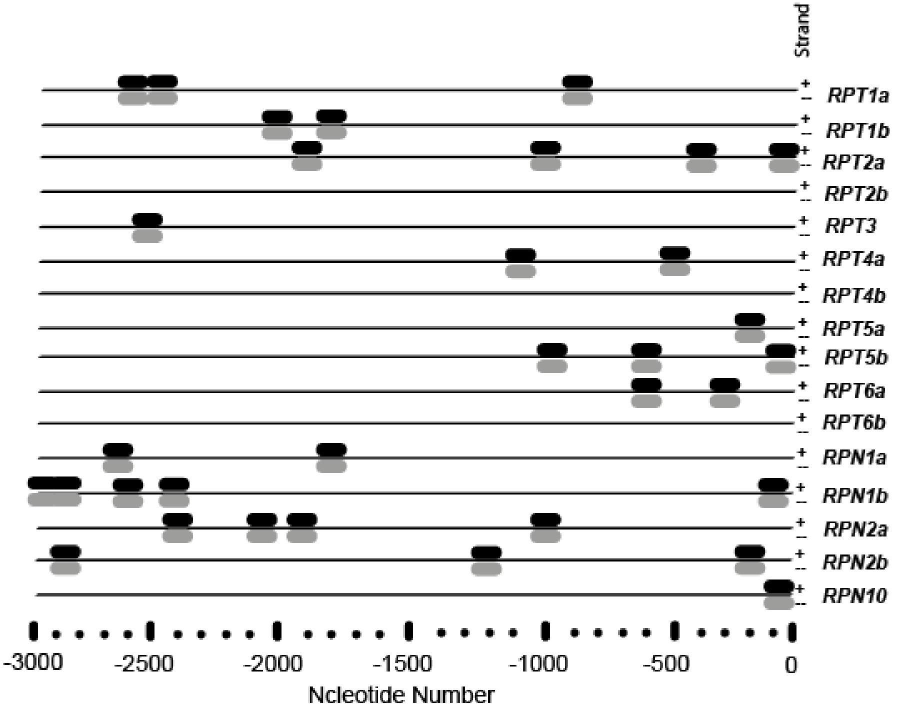
Strand
RPT1a
RPT1b
RPT2a
RPT2b
RPT3
RPT4a
RPT4b
RPT5a
RPT5b
RPT6a
RPT6b
RPN1a
RPN1b
RPN2a
RPN2b
RPN10
-3000
-2500
-2000
-1500
-1000
-500
0
Ncleotide Number

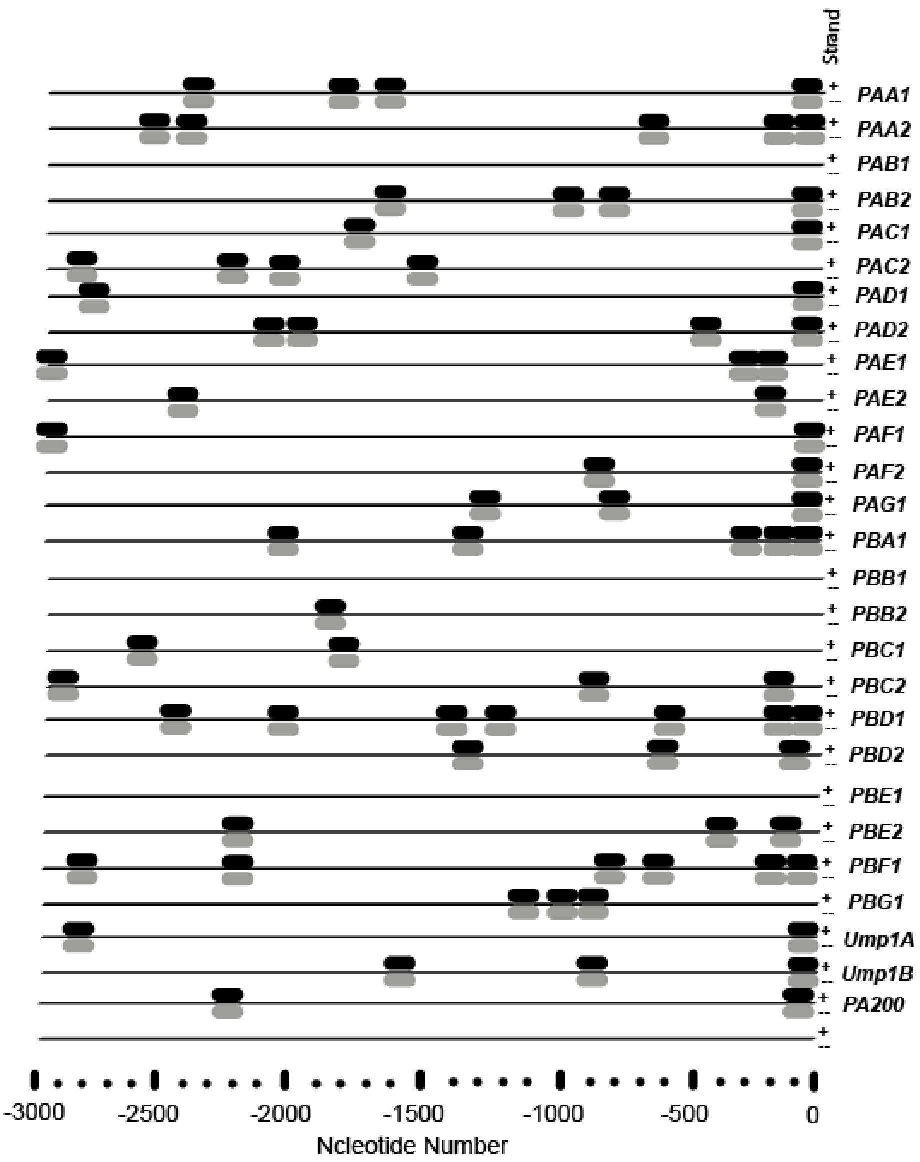


**Supplemental Figure S12.** NAC TFs recognition sequences in promoter regions of 26S proteasomal genes**.** Consensus sequence C[GT]TNNNNNNNA[AC]G was used as potential binding motif of NAC TFs to perform MEME analysis. Dark and gray boxes indicate the promoter position of palindromic sequences predicted by MEME analysis.

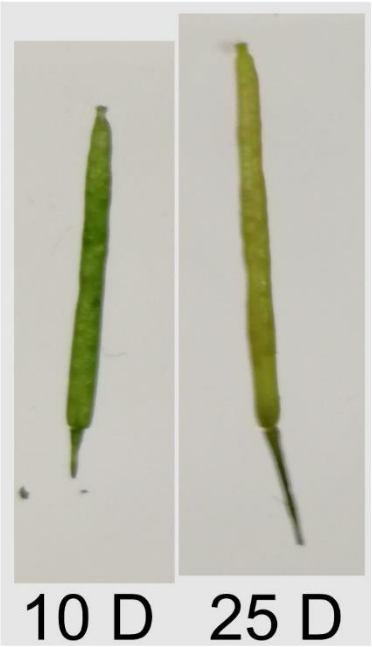


**Supplemental Figure S13.** Phenotype of collected siliques. Left a 10-day-old silique and on the right a 25-day-old silique, showing already clear signs of chlorophyll loss and senescence.
